# Cortical Layer 6b Persistent Subplate Neurons Reciprocally Connect Sensorimotor Areas and Inversely Reflect Somatosensory Engagement

**DOI:** 10.64898/2026.04.15.718699

**Authors:** Roksana Khalid, Débora Masini, Andrea Locarno, Olga Netsyk, Johan Lorentz, Christian Broberger

## Abstract

Persistent subplate neurons in layer 6b (L6b) of the cerebral cortex have long been assumed to be a passive remnant of early development. Recent findings are challenging this concept, but the functional role of L6b remains mysterious. A large group of L6b neurons express Connective Tissue Growth Factor (Ctgf). Here, we mapped L6b^Ctgf^ neurons throughout the cortex and found the highest density in primary sensorimotor areas. Cortical L6b^Ctgf^ projections are ipsilaterally restricted and origin-site specific, with notable reciprocity, whereas innervation of the thalamus is modality-matched. L6b^Ctgf^ neurons are electrically diverse and form excitatory synapses with both pyramidal cells and interneurons. In freely moving mice, these neurons are highly active and have prominent task-specific and temporally locked responses to external stimuli. These responses are dominated by paradoxical population silencing with escalating task engagement. Thus, L6b^Ctgf^ neurons may keep cortical ensembles in a state of alertness and available to process cognate stimuli.

**HIGHLIGHTS:**

- L6b^Ctgf^ neurons form origin-and layer-specific ipsilateral cortical connections.
- Electrically heterogeneous L6b^Ctgf^ neurons excite multiple classes of cortical neurons.
- Barrel cortex L6b^Ctgf^ neurons are highly active in freely moving mice.
- Somatosensory exploration paradoxically silences active barrel cortex L6b^Ctgf^ neurons.

## INTRODUCTION

The neocortex, the evolutionarily latest addition to the mammalian brain^1^, is instrumental in our experience of the world, our ability to integrate intero-and exteroceptive sensory experiences, and to select and execute appropriate actions. Attempts to understand the function and internal machinations of the cortical network have centered on the functional architecture of this stratified and regionally specialized structure, identifying cell types, connections, physiology and roles of the different layers^2,3^. Yet, until recently and with few exceptions^4–7^, scant attention has been paid to the neurons located deepest in the cortical mantle, the so-called persistent subplate neurons (PSNs)^8,9^.

This gap in our understanding of the cerebral cortex components is partly due to the assumption that PSNs are the passive remnants of a vestigial structure. Indeed, during the early formation of the neocortex, the origin of the PSNs, a structure known as the subplate^10^, plays an outsize role in its organization by guiding the projections and connections of cortical cells^10–12^ (see also^13^). Subsequently, most subplate neurons undergo a dramatic programmed cell death^8,14–16^. Yet, a smaller fraction of these cells – the PSNs – remains^9,16–19^ as scattered neurons in the white matter (WM neurons), and as a layer of cells (thinner in rodents, thicker in primates^20^) forming the cortical layer 6b (L6b^21^). These are separated from the more expansive L6a by a fibrous cell-poor leaflet^22,23^.

While long assumed to be functionally inert, research on PSNs in the past decade has challenged this view^24–32^. These studies have shown PSNs are anatomically and functionally integrated into the cortical microcircuit^28,29,33^, are sensitive to arousal neuromodulators^6,34–37^, exhibit axonal arborization patterns not elsewhere replicated among cortical projection neurons^27,38^, and able to elicit the electrophysiological^33^ and behavioral hallmarks of arousal^30,39^. Collectively, this work suggests that PSNs are important ‘citizens’ of the cortical population with unique functions.

PSNs are not a monolithic population, but rather encompass several subpopulations, as shown by the discrete, albeit partly overlapping, expression of molecular markers^40^. Thus, to understand the relevance of this “layer with no known function”^31^, the different PSN components need to be characterized with regard to cellular properties, connectivity and functional features. In previous work, we have explored these features in WM neurons^33,36^, where anatomical location aids identification. For the more molecularly heterogeneous L6b, we take advantage of the fact that the gene encoding Connective Tissue Growth Factor (Ctgf) is almost exclusively expressed in L6b within the adult CNS^28,40–43^. This pattern confers a specificity absent for several other L6b-expressed genes such as *e.g.* the dopamine D1 receptor, which can also be found in L6a^38,44^. Here, we have investigated the anatomical distribution, connectivity, intrinsic properties, local circuit interactions and behavior-correlated *in vivo* activity of L6b^Ctgf^ neurons. The results reveal structural and functional features that align with a role in task-associated arousal and sensory processing.

## RESULTS

### Ctgf-Cre mouse line shows high specificity to L6b neurons across cortical areas

In order to verify the validity of using the Ctgf-Cre mice as a platform to selectively study and manipulate L6b neurons, we first applied *in situ* hybridization to quantify *Ctgf* and *Cre* mRNA co-expression in these animals. All cortical neurons expressing *Ctgf* were tightly packed as a lamina of 80µm thickness abutting the dorsal border of the corpus callosum (distance measured for 783 *Ctgf* expressing neurons) and 97% also expressed *Cre* (903/933 neurons, primary somatosensory cortex, SSp; **Fig.1**).

**Figure 1.**
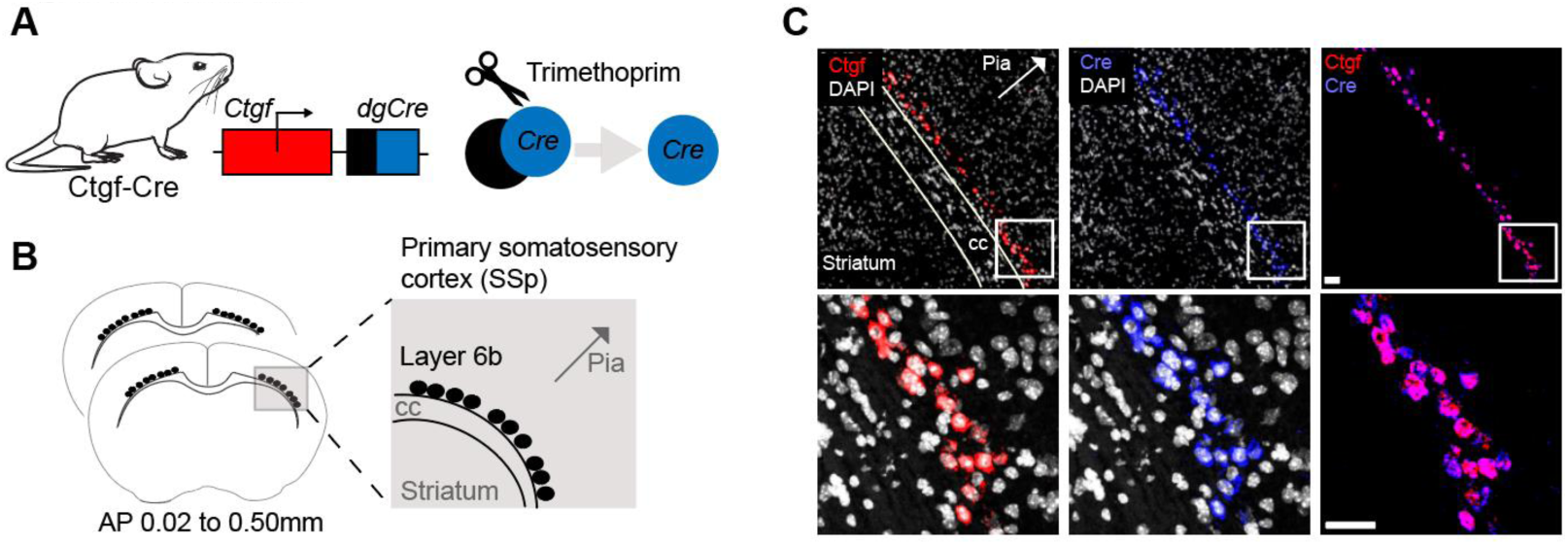
**Ctgf-Cre mice allow for selective labelling of cortical layer 6b neurons**. **(A)** Illustration of the Ctgf-Cre mouse line genetic construct which drives the expression of *Cre* under control of the *Ctgf* promoter. Functional Cre is dependent upon intraperitoneal injection of trimethoprim, thus Cre expression is under temporal control. **(B)** Schematics of the coronal sections used for *in situ* hybridization (ISH). **(C)** Confocal microphotographs depicting ISH for *Ctgf* and *Cre* mRNA and their co-expression in L6b of the SSp. *Ctgf* (red), *Cre* (blue), co-expression (magenta), nuclear DAPI (white). White boxes are magnified in the lower panel. 12 sections, six mice. Scale bar 50µm. cc, corpus callosum; Ctgf, Connective tissue growth factor; AP, antero-posterior position in relation to Bregma.

Next, we crossed Ctgf-Cre mice with a floxed-tdT reporter line in order to visualize *Ctgf* expressing neurons (tdT+) in the whole brain. We mapped the location of all tdT+ neurons across sagittal sections and registered their anatomical location to the Allen Mouse Brain Atlas (80µm/slice, one-in-four sampled, four mice). In total, 132 880 neurons were identified (sagittal sampling covered the mediolateral range from 3.70 to 0.00mm as per Paxinos^45^; total tissue area sampled: 69.4±8.8cm^2^).

Fluorescent cell bodies were present as a lamina juxtaposed to the WM, found across the entire cortical mantle, but showing higher density in the antero-medial cortex (**Fig.2A** and **Fig.S1A**). The highest cortical cell count occurred in the SSp (27% of all *Ctgf-*expressing cortical neurons) and primary motor area (MOp, 17% of all *Ctgf*^+^ cortical neurons; **Fig.2B**). The laminar thickness of L6b^Ctgf^ neurons was measured from the centroid of the most ventral to the centroid of the most dorsal neuron within each cortical region; we found layer thickness to vary by up to 67.6µm depending on the region evaluated. The MOp exhibited the greatest average L6b thickness (127.0±15.1µm, 30 levels sampled), whereas frontal/prefrontal areas (anterior cingulate and infralimbic) had narrow bands (71.0±3.3µm, 18 levels sampled; **Fig.2C**). Only 3.45% of all *Ctgf*-expressing neurons were located outside the neocortex (4 589 neurons, **Fig.2D**). Within this minority population, the most prominent non-cortical *Ctgf*-expressing cell clusters were in olfactory areas (including the main and accessory olfactory bulbs), followed by the endopiriform nucleus and hippocampal region (**Fig.2E**). Given this anatomical distribution, and in agreement with earlier reports^29,46,47^, we conclude that the Ctgf-Cre mouse line offers a tractable platform for the study of L6b neurons with laminar specificity across cortical areas. The observed enrichment of *Ctgf*-expressing neurons in primary sensorimotor cortex parallels reports in macaque and human neocortex^16^ describing regionally variable survival of PSNs, with particularly high densities in primary areas, indicating a conserved organizational principle.

**Figure 2.**
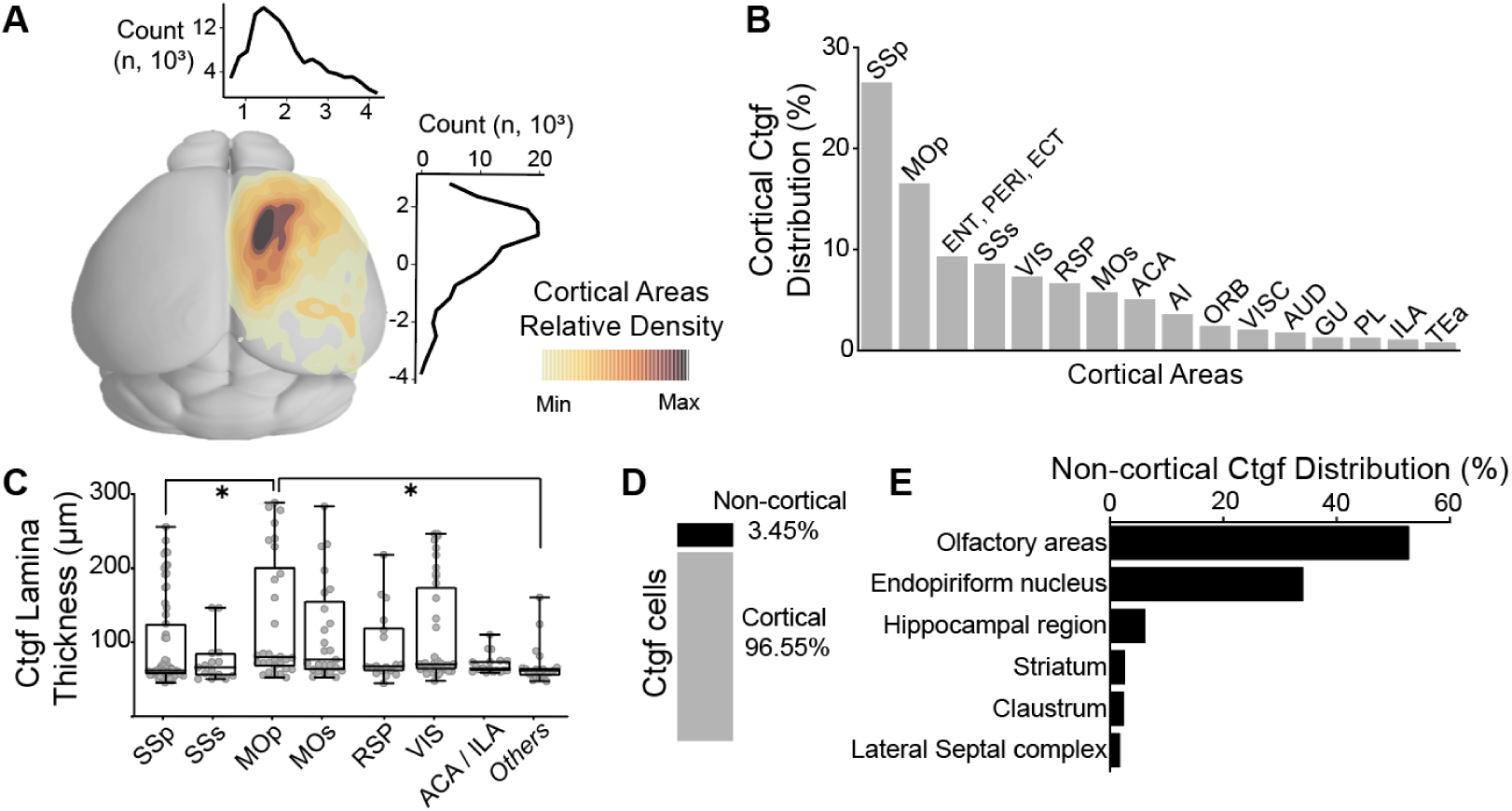
L6b^Ctgf^ neurons are more densely packed in the SSp and MOp than in other cortical areas. **(A)** Cortical Ctgf cell density with quantification along the anteroposterior and mediolateral axes (four mice). **(B)** Percentage of Ctgf neurons found in L6b across different cortical areas over the total number of cortical Ctgf cells (128 291 cortical Ctgf neurons mapped, four mice). **(C)** Lamina thickness estimation of L6b based on Ctgf neuron distance from the white matter. The class named ‘Others’ includes areas of the insular cortex (GU, VISC, AI, TEa). Kruskal-Wallis H(8)=23.07, p=0.0017, Dunn’s multiple comparisons, all *vs.* all; Others *vs.* MOp p=0.0114, SSp *vs.* MOp p=0.0290, all else p>0.05. Tukey min-max bars. **(D)** Proportion of Ctgf neurons in cortical and non-cortical areas (4 589 non-cortical neurons mapped, four mice). **(E)** Distribution of non-cortical Ctgf neurons. Of all non-cortical Ctgf cells (percent/region); 53% olfactory areas, 34% endopiriform nucleus and 6% hippocampal region. No other area accounted for more than 3% of the total extracortical cell count. ACA, Anterior cingulate area. AI, Agranular insular cortex. AUD, Auditory cortex. ECT, Ectorhinal cortex. ENT, Entorhinal cortex. GU, Gustatory cortex. ILA, Infralimbic cortex. MOp, Primary motor cortex. MOs, Secondary motor cortex. ORB, Orbital cortex. PERI, Perirhinal cortex. PL, Prelimbic cortex. RSP, Retrosplenial cortex. SSp, Primary somatosensory area. SSs, Supplemental somatosensory area. TEa, Temporal association cortex. VISC, Visceral cortex. VIS, Visual cortex. See also **Figure S1A**.

### The axonal arbors of primary somatosensory L6b^Ctgf^ neurons are ipsilaterally restricted and preferentially target the motor cortex

Dense intracortical axons emanating from L6b have been described in cats^48^, ferrets^49^ and rodents^24,50^. To map the axonal arbors of anatomically defined L6b^Ctgf^ neurons, we injected a Cre-dependent virus that induces the expression of two physically separated color tags^51^, a membrane-tethered enhanced green fluorescent protein (EGFP) and a red fluorescent protein (mRuby) linked to murine synaptophysin, a marker of synaptic boutons (DIO-EGFP-syp1-mRuby)^52^. This approach allows for a comprehensive analysis of L6b^Ctgf^ axonal collateralization (or lack thereof). Since we identified the SSp as the cortical area with the highest density of Ctgf labeled neurons, we targeted this region with a unilateral viral injection. Following a four-week incubation period, we mapped projection sites using serial imaging of the entire brain (coronal sections, four mice with total tissue area surveyed of 17.3cm^2^; sagittal sections, two mice) (**Fig.3A**).

**Figure 3.**
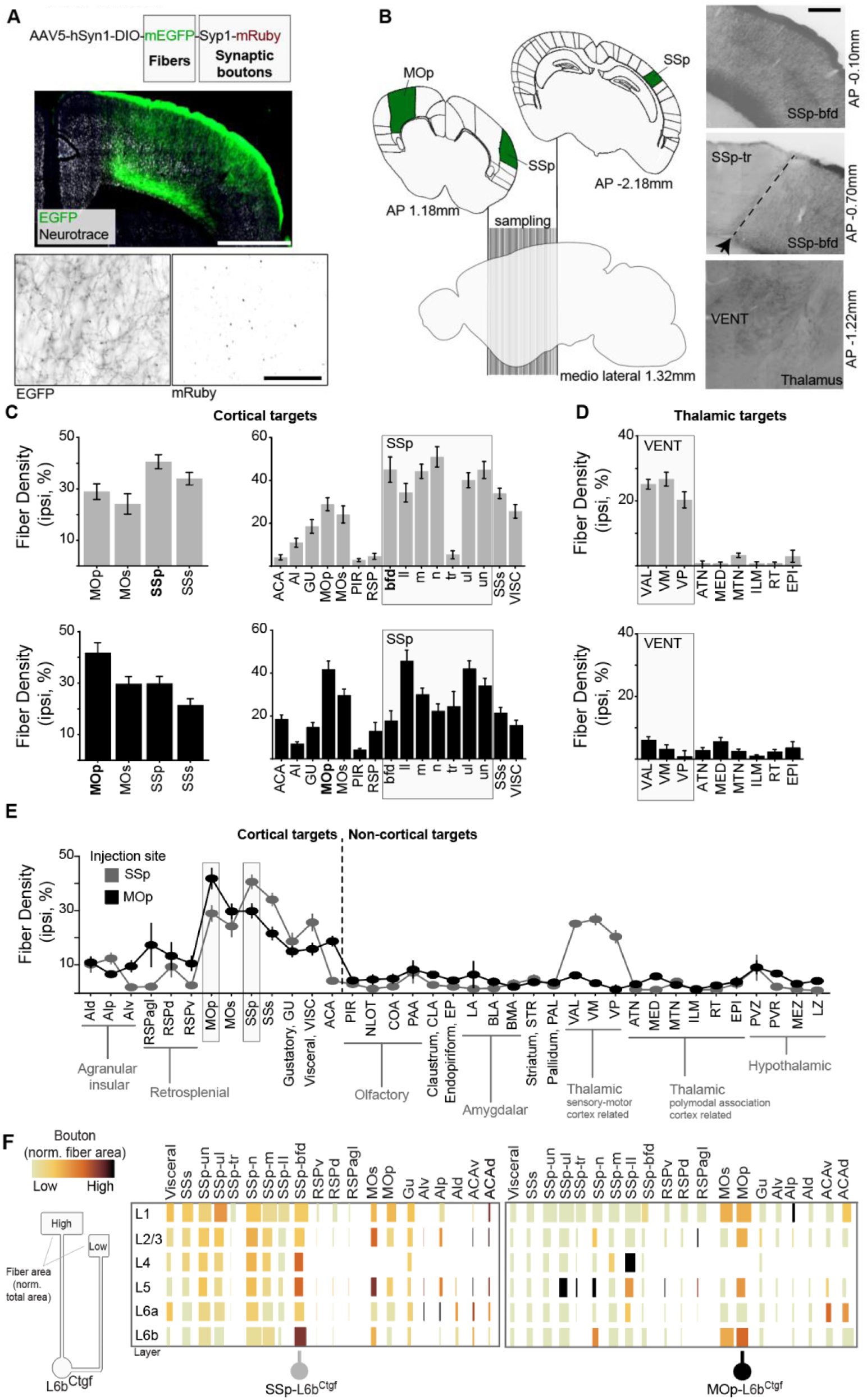
L6b^Ctgf^ axons are ipsilaterally restricted, form dense intracortical connectivity, and show preferential SSp-MOp reciprocity. **(A)** Viral tracing of L6b^Ctgf^ axons and synaptic boutons. Confocal images of wide view (scale bar 1mm) and a closeup showing separated channels for fibers (EGFP) and synaptic boutons (mRuby) (black and white, inverted; lower row, scale bar 50μm). **(B)** Left, schematic representation of the coronal slices used for mapping L6b^Ctgf^ connectivity (two groups; SSp injected and MOp injected, highlighted in green). Sections were collected from anteroposterior 1.18 to-2.18mm. Right, representative images from a SSp injected mouse showing L6b^Ctgf^ fibers in the SSp-bfd (top), SSp-tr/SSP-bfd border (middle), and VENT (bottom). Scale bar 200μm. **(C)** L6b^Ctgf^ fiber density across cortical areas, from SSp-L6b^Ctgf^ (top) and MOp-L6b^Ctgf^ (bottom) injected mice. Left, subregions belonging to the same cortical region are averaged together. Right, cortical subregions are specified. **(D)** Fiber density in thalamic targets. SSp-L6b^Ctgf^ (top) and MOp-L6b^Ctgf^ (bottom) injected mice. **(E)** Fiber densities for all brain regions analyzed. SSp-L6b^Ctgf^ (gray) and MOp-L6b^Ctgf^ (black) injected mice. **(F)** For each cortical layer, fiber density (box size) and bouton density (heatmap) normalized to total area. SSp-L6b^Ctgf^ (left) and MOp-L6b^Ctgf^ (right) injected mice. MOp, Primary motor cortex. SSp-bfd, Primary somatosensory barrel field. SSp-tr, Primary somatosensory trunk. VENT, Ventral group of the thalamus. A complete list for anatomic acronyms for this figure can be seen in **Table S4** or at the Allen Atlas^53^ webpage. See also **Figure S1B-E**.

SSp-L6b^Ctgf^ neurons exhibited axonal arbors that expanded rostrally as far as the cortical frontal pole (2.40mm rostral to injection site) and as far caudally as the visual cortex (3.50mm caudal to injection site). These long-range projections were ipsilaterally restricted and traveled across cortical areas while remaining within the boundaries of L6 and largely avoiding crossing through the WM, which is the primary path for long-range projection cortical axons (**Fig.S1B-C**). Once axons arrived at a target region, they projected towards the pia mater in nearly perpendicular paths which precisely marked the borders between cortical regions (**Fig.3B**). The density of L6b^Ctgf^ fibers was highest at the injection site (average fiber coverage SSp, 41±3%, 43 levels sampled; SSs, 34±2%, 43 levels sampled) and the main target area was the MOp (29±3%, 41 levels sampled) (**Fig.3C**, gray bars). Notably, SSp-L6b^Ctgf^ fibers avoided SSp-trunk (SSp-tr), a region positioned between the injection site (SSp) and the main target MOp (**Fig.3B**). Hence, the observed SSp-to-MOp connectivity reflects genuine pathway specificity rather than inadvertent injection spread.

Within each cortical subdivision, axonal L6b^Ctgf^ collateralization occurred at specific laminae. For example, fibers targeting the MOp had the highest densities in L1 (38±4%) and L2/3 (28±3%). In stark contrast, the dorsal agranular insular cortex (AId) was completely devoid of detectable fibers in L1 with axons stopping at L5. Hence, SSp-L6b^Ctgf^ axonal distribution within the cortex is not only regionally distinct but also layer specific.

SSp-L6b^Ctgf^ axons also innervated deep, subcortical structures. They were seen to occasionally cross the WM and enter the dorsal striatum, then travel ventro-caudally towards the globus pallidus, after which they would cross the corticospinal tract as bundles and move medially via the reticular nucleus of the thalamus (without making synaptic contacts in this structure) to finally terminate within thalamic areas (ventral group of the thalamus, VENT which is a thalamic sensory-motor cortex related area; fiber coverage was between 20-27%; **Fig.3D**, gray bars). No SSp-L6b^Ctgf^ axons reached the hippocampal area, nor was there significant innervation of other subcortical regions, such as the claustrum, endopiriform nucleus or amygdalar areas (**Fig.3E**).

To locate SSp-L6b^Ctgf^ synaptic boutons and differentiate between true target areas and fibers of passage, we analyzed the number of mRuby-puncta. Values were normalized to fiber density and target area size. Within the SSp, the highest relative puncta density was present within the L6b, followed in sequence by L1>L4/L5 (**Fig.3F**). Even though innervation of the MOp was higher, the secondary motor region (MOs) was more puncta-dense with SSp-L6b^Ctgf^ axons forming synaptic boutons preferentially to MOs-L5 and MOs-L2/3 (**Fig.3F**, left panel). Projections towards the sensory-motor related area of the thalamus (*i.e.,* VENT), which equally innervated all its subregions, had strikingly variable puncta density. The highest density was recorded in the ventral-posterior complex (VP). In contrast, puncta density in the anterior-lateral complex (VAL) was approximately 50% of that in VP (VAL=0.51xVP). Finally, the medial nucleus (VM) had only sparsely located synaptic boutons (VM=0.11xVP, 89% less than VP). Therefore, within the VENT, the VP receives the highest bouton density and thus represents the primary subcortical synaptic target of SSp-L6b^Ctgf^ projections (**Fig.S1D-E**). These data also highlight that caution is warranted when using fiber density as a direct proxy for synaptic targeting.

### Somatosensory and motor areas share reciprocal ipsilateral connections via L6b^Ctgf^ neurons

Having identified that SSp-L6b^Ctgf^ neurons preferentially project towards, and form synaptic boutons with, the ipsilateral motor cortex, we next examined if the MOp reciprocates this connection, by injecting the MOp with the same viral construct (coronal sections, four mice with 22.48cm^2^ tissue area surveyed; sagittal sections, two mice).

MOp-L6b^Ctgf^ axonal arbors were also ipsilaterally restricted and the largest part of MOp-L6b^Ctgf^ fibers remained within the injected area (average 42±4%, highest density within L1, 63±4%) whereas MOs and SSs were targeted without a clear preference (MOs 30±3%, SSs 21±3%; **Fig.3C**, black bars). Notably, the SSp was the main target area outside the motor cortex (SSp 30±3%). Whereas thalamic targets exhibited negligible innervation (**Fig.3D-E** and **Fig.S1E**). MOp-L6b^Ctgf^ neurons had their highest puncta count in the MOs, within L1 and L6b. Once again, bordering somatosensory areas, had distinct and layer dependent puncta densities (**Fig.3F**, right panel).

These findings show that L6b^Ctgf^ neurons in the SSp and MOp establish reciprocal, ipsilateral corticocortical connectivity characterized by origin-specific projection patterns, distinct regional and laminar targeting, and a shared bias toward sensorimotor-related regions.

### L6b^Ctgf^ neurons form glutamatergic monosynaptic connections with both principal neurons and interneurons of the cortex

To explore the properties of L6b^Ctgf^ connections to their local targets, we injected a Cre-dependent virus encoding Channelrhodopsin-2 (ChR2, mCherry tag) unilaterally in either the SSp or the MOp. Whole-cell patch-clamp recordings in acute coronal slices confirmed that L6b^Ctgf^ neurons expressing ChR2 were viable and responded to photo-stimulation (10ms pulses, 470nm). Spike fidelity was maintained for up to 20Hz (**Fig.4A-B**).

**Figure 4.**
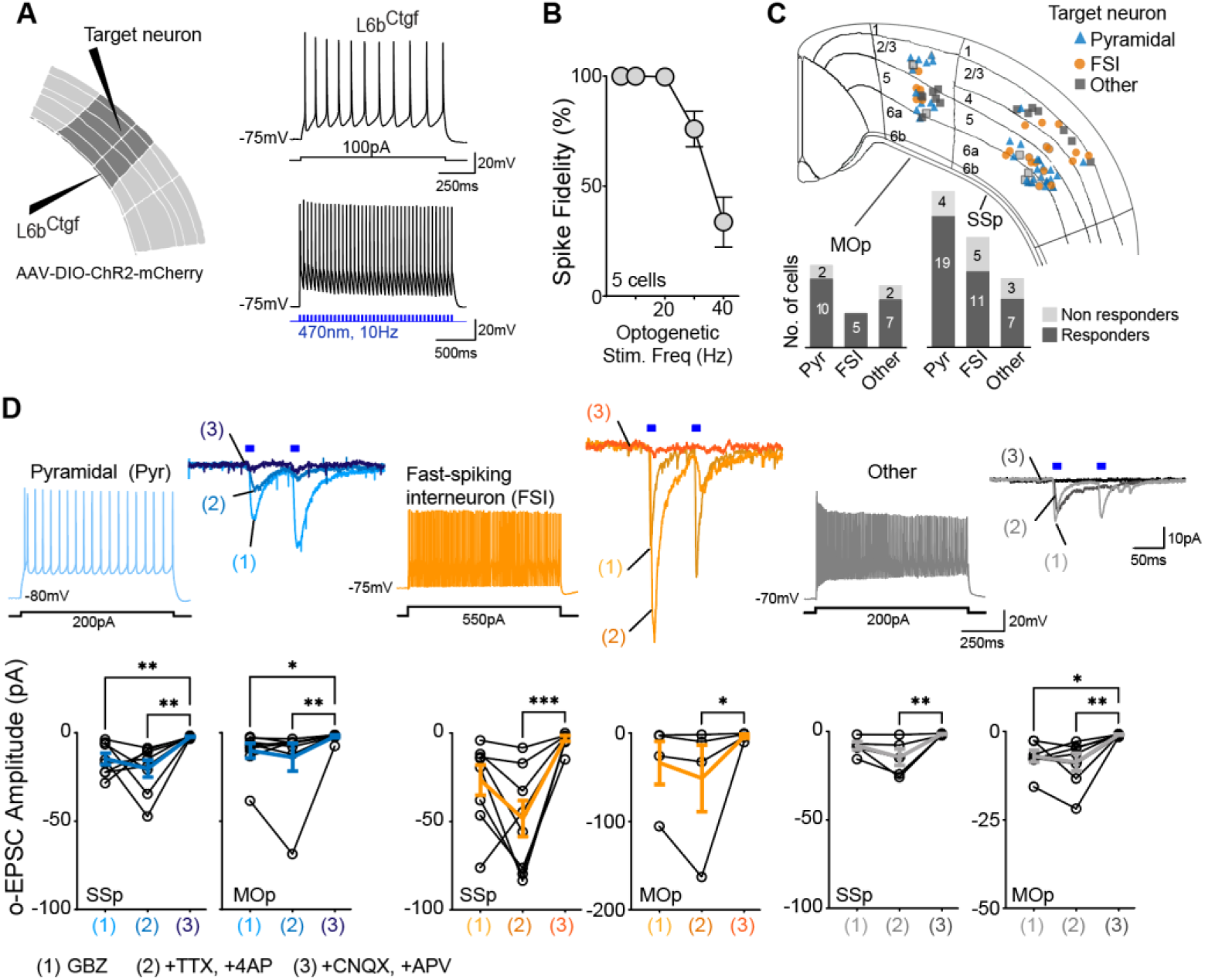
L6b^Ctgf^ neurons synapse onto principal neurons and interneurons within the cortex. **(A)** Left, schematic of optogenetic experiments. Either SSp or MOp were transduced with a Cre dependent ChR2 expressing virus. Right upper panel shows a patched L6b^Ctgf^ neuron spiking in response to 100pA depolarizing square pulse; lower panel shows the same neuron responding to 10Hz optogenetic activation (blue). **(B)** Quantification of spike fidelity at different optogenetic stimulation frequencies. Spike fidelity was high for frequencies ≤20Hz (five cells from three mice). **(C)** Localization and cell identity of recorded neurons across cortical layers. Bar graphs, number of responder and non-responder neurons in SSp and MOp, grouped by cell type, in response to optogenetic activation of surrounding L6b^Ctgf^ axons. Responded to optogenetic stimulation: 83% (29/35) of pyramidal neurons, 76% (16/21) of FSI neurons and 74% (14/19) of interneurons classified as ‘Other’. Total of 75 cells from 14 mice, 2-5 slices/mouse. **(D)** Top, electrophysiological signatures of target neurons in response to current injection, used for neuronal type classification as pyramidal (Pyr, blue), FSI (orange), or ‘Other’ (gray). Next, L6b^Ctgf^ terminals are opto-activated twice in rapid succession (blue rectangles), and the resultant paired-pulse o-EPSC responses are measured during sequential application of synaptic blockers: stage 1, GBZ is added to the bath, followed by (stage 2) TTX and 4AP, and finally (stage 3) CNQX and APV. Lower panel, o-EPSC response quantification showing that response to optogenetic L6b^Ctgf^ activation is lost only when monosynaptic glutamatergic connections are blocked. Stats for SSp: Friedman test repeated measurements. Pyr, Q=12, p=0.001. FSI, Q=14, p<0.0001. Other, Q=10, p=0.0008. Dunn’s multiple comparisons, all *vs.* all. Stage 1 *vs.* 2, all p>0.05. Stage 1 *vs.* 3, Pyr p=0.008; FSI p=0.07, Other p=0.3415. Stage 2 *vs.* 3, Pyr p=0.008; FSI p=0.0005, Other p=0.005. Stats for MOp: Friedman test repeated measurements. Pyr, Q=14, p=0.0009. FSI, Q=7, p=0.04. Other, Q=11, p=0.003. Dunn’s multiple comparisons, all *vs.* all. Stage 1 *vs.* 2, all p>0.05. Stage 1 *vs.* 3, Pyr p=0.02; FSI p= 0.23, Other p=0.02. Stage 2 *vs.* 3, Pyr p=0.004; FSI p=0.04, Other p=0.01. SSp, primary somatosensory cortex. MOp, primary motor cortex. Drug related acronyms: 4AP, aminopyridine. APV, D-2-amino-5-phosphonopentanoate. CNQX, cyanquixaline. GBZ, gabazine. TTX, tetrodotoxin. See also **Figure S2A-B** for description of paired pulse facilitation and **Table S3** for statistical details.

We tested if optogenetic activation of L6b^Ctgf^ axons could elicit synaptic responses in other cortical neurons, using localized LED illumination (317µm around the patched cell; synaptic response peak latency <10ms from light onset). Patched neurons were identified as pyramidal (35 cells) or fast-spiking interneurons (21 cells) based on their electrophysiological response to depolarizing current pulses^54–57^. All remaining putative interneurons (19 cells) were classified as ‘Other’. Light-evoked excitatory postsynaptic currents (o-EPSCs) were isolated by bath application of the GABA_A_ receptor antagonist gabazine (GBZ, 10µM), and recorded in voltage clamp mode at-70mV. We identified L6b^Ctgf^ evoked o-EPSCs in the majority of the recorded neurons within both SSp (75.51%, 49 cells) and MOp (84.62%, 26 cells) (**Fig.4C**).

L6b^Ctgf^ triggered o-EPSCs likely represent monosynaptic connections, as they persisted when we blocked polysynaptic responses using TTX (1µM) and 4-aminopyridine (4AP; 100µM), abolishing voltage-gated Na^+^ and K^+^ currents, respectively. Furthermore, the evoked o-EPSCs were mediated by ionotropic glutamate receptors, as they were abolished by application of cyanquixaline (CNQX, 10µM) and D-2-amino-5-phosphonopentanoate (APV, 10µM). Monosynaptic glutamatergic responses were observed in both SSp and MOp (**Fig.4D**).

To evaluate the strength of these excitatory synapses, we calculated paired-pulse ratio as a measure of glutamate release probability^58^. Facilitation (larger amplitude responses to the second light-pulse) was present in all neuronal types (**Fig.S2A-B**). Responses by pyramidal neurons were also evaluated layer-wise and no differences in absolute amplitudes were found (Kruskal-Wallis H(3)=2.27, p=0.52; **Table S3**). These data demonstrate that L6b^Ctgf^ neurons form extensive, translaminar connectivity via facilitating glutamatergic synapses with diverse neuronal types.

### Neighboring L6b^Ctgf^ neurons are not monosynaptically interconnected

Next, to identify potential functional interconnectivity among L6b^Ctgf^ neurons, we performed paired patch-clamp recordings (14 neurons, five mice). Injection of depolarizing square-pulse current steps in one neuron reliably elicited action potentials, but failed to evoke any postsynaptic response in the neighboring L6b^Ctgf^ neuron (**Fig.S3A-C**[ii-iii], and **E**). Likewise, hyperpolarizing current injections did not elicit correlated voltage deflections in the paired neuron (**Fig.S3C**[iv-v] and **F**). These data indicate the absence of detectable chemical or electrical synaptic transmission between neighboring L6b^Ctgf^ pairs, arguing against impactful local interconnectivity within the population.

Although no direct synaptic coupling was detected, we occasionally observed simultaneous spontaneous postsynaptic potentials (PSPs) in paired recorded L6b^Ctgf^ neurons (**Fig.S3C**[i]). To assess whether these coincident events, indicating synchronous input activity, exceeded chance levels, we computed the cross-correlation of all spontaneous PSPs between pairs. This analysis revealed a peak centered around zero lag (peak amplitude: 2.9±0.3 z-score; lag: 4.6±1.2ms; **Fig.S3D**), showing that coincident spontaneous PSPs occurred more frequently than expected by chance. These results are consistent with weak, but non-random, temporal coordination between L6b^Ctgf^ neurons, likely reflecting co-activation by shared presynaptic inputs, but not direct synaptic or electrical interconnectivity.

### The firing patterns and membrane properties of L6b^Ctgf^ neurons are heterogeneous and comparable in diversity to those of neighboring L6b neurons

The intrinsic firing properties of L6b^Ctgf^ neurons were evaluated through perforated patch-clamp electrophysiology (33 cells, 27 mice), to optimize membrane integrity and minimize dialysis of the cytoplasm^59^ (**Fig.5A**). For comparison, we also recorded L6b non-*Ctgf* cells (L6b^Others^; non-fluorescent). L6b^Others^ neurons (18 cells, six mice) were selected based on their distance to the WM (within 50µm) and their proximity to a fluorescent L6b^Ctgf^ neuron (average distance, 26±7µm).

**Figure 5.**
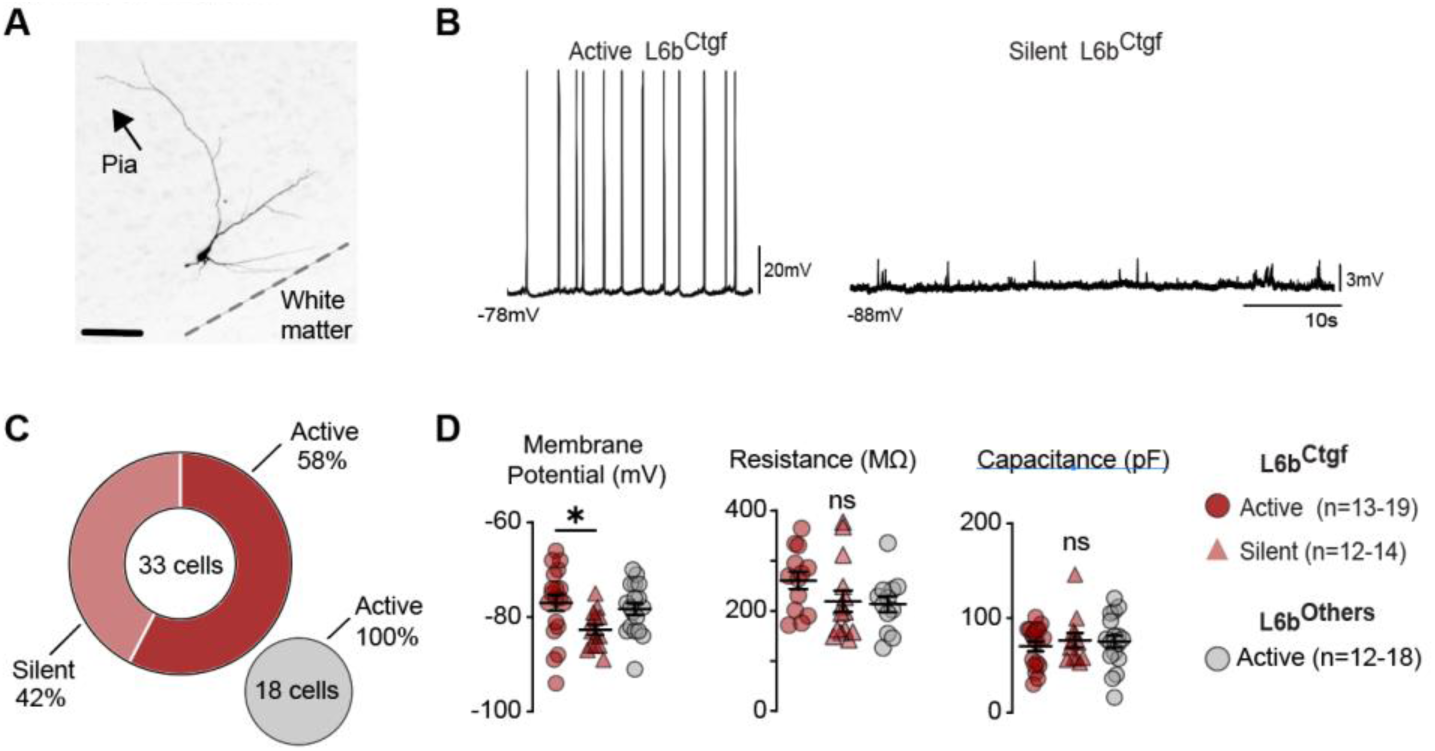
Activity patterns and passive membrane properties of L6b^Ctgf^ and neighboring L6b neurons. **(A)** Micrograph of a neurobiotin filled SSp-L6b^Ctgf^ neuron. Scale bar 100µm. SSp, primary somatosensory cortex. **(B)** Representative traces from active (left) and silent (right) L6b^Ctgf^ neurons recorded in perforated patch current clamp configuration. Note difference in membrane potential. **(C)** Proportion of active *vs.* silent L6b^Ctgf^ (red) during baseline recording. Neighboring L6b neurons, not expressing the tdTomato marker, were all active during baseline recording (gray, L6b^Others^). L6b^Ctgf^ 33 cells from 27 mice. L6b^Others^ 18 cells from six mice. **(D)** Quantification of L6b neuronal passive properties. L6b^Ctgf^ active neurons were significantly more depolarized than the silent subset. Input resistance and capacitance were similar between groups. Recordings performed in the SSp. Stats for Membrane potential: one-way ANOVA F=3.8, p=0.03. Tukeýs *post hoc* test: L6b^Ctgf^ active *vs.* L6b^Ctgf^ silent p=0.03; L6b^Ctgf^ active *vs.* L6b^Other^ p=0.78; L6b^Ctgf^ silent *vs.* L6b^Others^ p=0.11. Mean±SEM (mV), L6b^Ctgf^ active-77.0±1.74, L6b^Ctgf^ silent-82.7±1.07, L6b^Others^-78.3±1.27. Stats for Input resistance: Kruskal-Wallis H(2)=4.1, p=0.13. Mean±SEM (MΩ), L6b^Ctgf^ active 260.5±17.5, L6b^Ctgf^ silent 219.2±21.3, L6b^Others^ 213.2±16.1. Stats for Capacitance: Kruskal-Wallis H(2)=0.2, p=0.89. Mean±SEM (pF), L6b^Ctgf^ active-77.0±1.74, L6b^Ctgf^ silent-82.2±1.07, L6b^Others^-78.3±1.27. See also **Figure S2C-G** for replication and validation experiments using the whole-cell patch-clamp method. **Figure S2H-J** for multispike activity. **Table S3** for statistical details.

Neurons were classified as either active or silent based on the presence of APs during baseline recordings with active neurons discharging in a wide range of spontaneous frequencies (0.01 to 12.40Hz; **Fig.5B**). Whereas all L6b^Others^ were active (18/18 cells), nearly half of L6b^Ctgf^ neurons were silent (42%, 14/33 cells; **Fig.5C**). Silent L6b^Ctgf^ neurons were more hyperpolarized than the active set, yet neither input resistance, nor capacitance, differed between groups (**Fig.5D**). We validated these results using whole-cell patch-clamp electrophysiology and found a similar proportion of active *vs.* silent L6b neurons as well as intrinsic properties (**Fig.S2C-G**).

To evaluate firing patterns, we extracted the interspike interval between APs and found a bimodal right-skewed distribution (**Fig.S2H**), indicating the presence of transient episodes of high frequency spiking^60^ (multispike events; **Fig.S2I**). Hence, active neurons can be either tonic firing or multispiking. Among multispiking neurons, no significant difference in the average number of spikes in a train was found between L6b^Ctgf^ and L6b^Others^ (**Fig.S2J**).

We also evaluated the responses of L6b^Ctgf^ (22-26 cells) and L6b^Others^ (11-12 cells) to depolarizing and hyperpolarizing current steps. Depolarizing currents of at least 15pA triggered AP discharge in both groups (Rheobase: L6b^Ctgf^ 37.5±4.5pA, L6b^Others^ 30.9±3.2pA; p=0.82, Mann-Whitney) with similar firing threshold, peak amplitude, peak-to-peak frequency and max rise slope (**Fig.S4A-C**). At the end of the depolarizing current step, neurons produced an after-hyperpolarization potential (AHP) with no differences in amplitude or timing (**Fig.S4D**). Finally, in response to hyperpolarizing steps, 72.9±6.2% of all neurons showed a sag potential (L6b^Ctgf^ 19/24 cells, L6b^Others^ 8/12 cells) without amplitude differences between groups, and both groups showed a post-pulse rebound potential of similar amplitude (**Fig.S4E**).

Thus, although several parameters were reviewed, we did not identify electrophysiological characteristics that consistently distinguish L6b^Ctgf^ neurons from neighboring L6b neurons. Instead, we found comparable electrophysiological diversity, suggesting that *Ctgf* expression serves primarily as a molecular marker of L6b rather than defining a phenotypically distinct neuronal subtype within it.

### L6b^Ctgf^ neurons exhibit non-recurrent, population-sparse activity in awake freely behaving mice

Our anatomical tracing showed that L6b^Ctgf^ neurons exhibit strong origin-specific long-range projections with distinct and region-selective targets. This organization suggests L6b^Ctgf^ neurons may be tuned to sensory cues and motor states specific to the cortical region in which they reside. To test this possibility, we focused on L6b^Ctgf^ neurons of the SSp-bfd since this region supports an active scanning somatosensory system^61^. To examine L6b^Ctgf^ activity, at single-neuron resolution, we expressed GCaMP8m and implanted a prism lens. We imaged L6b^Ctgf^ calcium fluctuations in freely moving mice as they engaged in spontaneous, self-initiated behaviors (four mice in all behavioral sessions, two males and two females; **Fig.6A-B**). Across six behavioral sessions we identified a total of 255 L6b^Ctgf^ neurons. Neuronal activity exhibited a wide dynamic range with per-cell signal variance spanning from 36 to 1.26×10⁵ (ΔF/F), confirming that all included neurons displayed detectable signal fluctuations.

**Figure 6.**
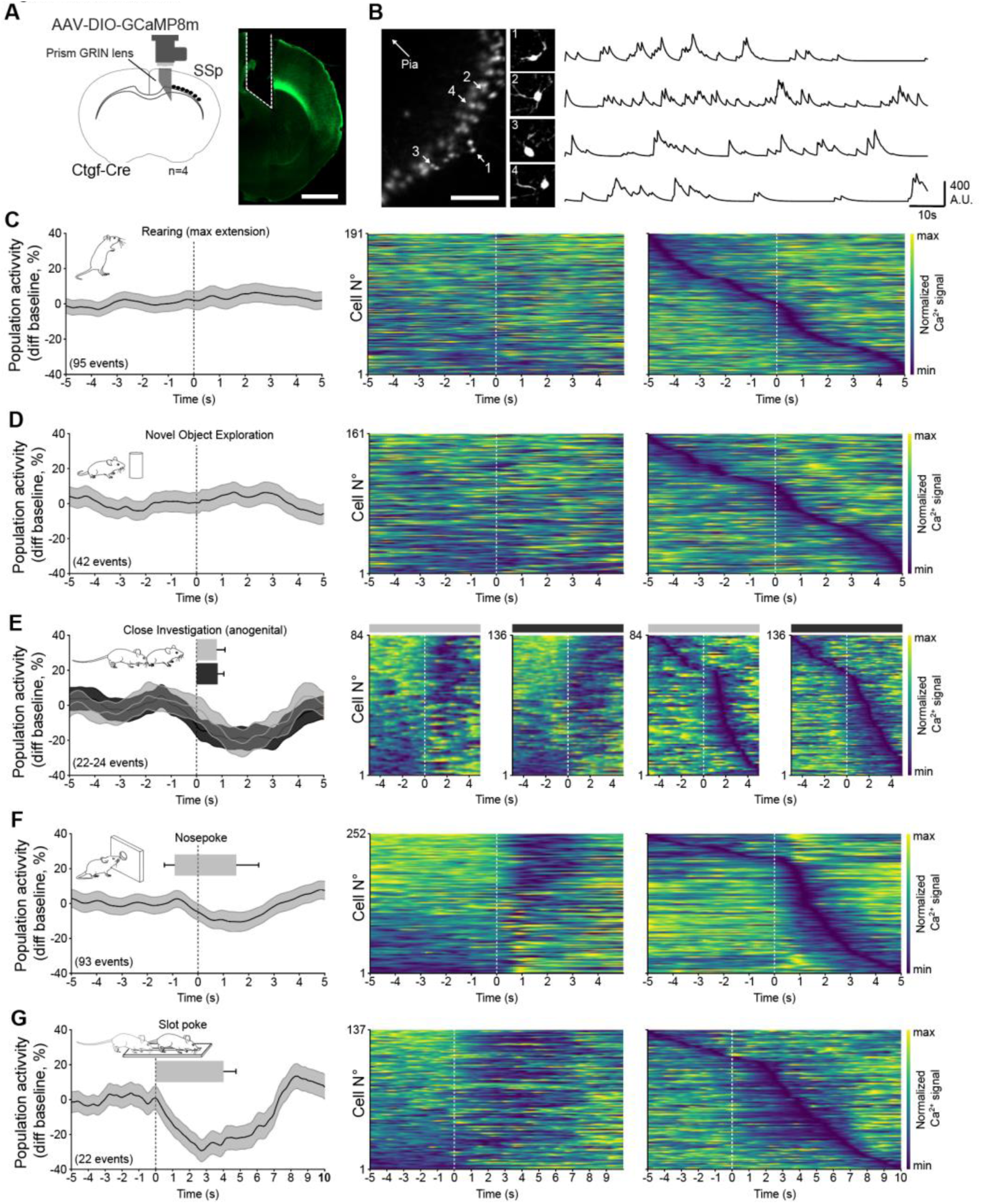
In freely moving mice, L6b^Ctgf^ neuronal activity is modulated by the external environment and time locked to behavior. **(A)** Left, schematic showing the miniscope position and viral strategy used to visualize L6b^Ctgf^ neuron activity within the SSp, subregion barrel field (four mice, scale bar 1mm). Right, microphotograph showing GRIN-prism lens placement. GCaMP8m-expressing Ctgf neurons are seen in green. **(B)** Miniscope field-of-view and example neurons (max projection image) with their respective activity traces. Scale bar 100µm. **(C-G)** Left, grand average fluorescence population signal (mean±SEM), around the annotated behavioral timepoint (vertical dashed line). Horizontal bars are plotted for behaviors in which bout duration was annotated. Heatmaps showing normalized cell activity averaged across behavioral events, organized by activity level (most active on top, middle) or time of trough (right). **(C)** Rearing. Zero marks moment of maximal vertical extension. **(D)** Object exploration. Zero marks maximal proximity. **(E)** Social close-investigation, anogenital sniffing. Gray, same sex. Black, opposite sex. Zero marks initiation of contact. **(F)** Nosepoke. Zero marks maximal nostril insertion into the poke-port. **(G)** Slot poke. Zero marks start of exploration. GRIN lens, Gradient Refractive Index lens. SSp-bfd; Somatosensory primary cortex, barrel field. See also **Figure 7** for a by class response analysis and **Figure S5**.

First, mice freely explored an empty open-field arena for 20 minutes while L6b^Ctgf^ neurons were continuously recorded (**Fig.S5A**). Temporal autocorrelation was computed to assess whether individual neurons displayed recurring activity patterns. The autocorrelation peak of each neuron was compared against a block-bootstrapped null distribution (5s window). Only 6.21% of neurons (9/145, false discovery rate [FDR] correction, q<0.01) exhibited recurrent activity with a peak amplitude of 0.24±0.02 (range: 0.16 to 0.35) and an associated frequency of 0.0031±0.0003Hz (range: 0.0023 to 0.0055Hz). We concluded that recurring neuronal activity is restricted to a minority of SSp-bfd L6b^Ctgf^ neurons in freely moving mice.

Next, we quantified sustained activation episodes to capture activity states other than periodic recurrence. Across animals, neurons exhibited on average 121 sustained activation episodes per session (range: 37 to 236 episodes for any given neuron/session) with a stereotyped active state duration across neurons and animals of 1.06±0.10s (range: 0.84 to 1.11s). Furthermore, for any given 500ms window, an average of 15.88±0.46% of L6b^Ctgf^ neurons were concomitantly active. At peak periods, up to 79.72±5.38% of all neurons engaged. Yet, such widespread activation was rare, in a 20min session, periods with >70% of neurons simultaneously active amounted to only 2.77±1.46s (cumulative average of animals). Therefore, during open-field exploration, a fraction (approx. 15%) of L6b^Ctgf^ neurons is always active, but large-scale co-activation events occur only rarely and for very short periods.

The observation that a subset of neurons is always active raises the possibility that the same cells tend to co-engage over time, which would manifest as consistent co-fluctuations between neuronal pairs. Across the 3 355 possible pairs, 316 (9.42%) showed significant co-fluctuation (pair circular-shift control, 10^3^ iterations per pair and FDR correction, q<0.01) approximately 4-fold larger than chance (population circular-shift, 10^3^ iterations per animal, null hypothesis produced 75.7 pairs, CI: 68 to 83). In co-fluctuating pairs, mutual information (MI) confirmed that activity in one neuron provides information about the other’s activity state (MI mean value across animals, range: 0.41 to 0.51), indicating these pairs have a temporal coupling which is more frequent than expected by chance, and that their pairing is not deterministic (MI<<1). Interestingly, the majority of pairs (281/316, 88.92%) showed bi-directional co-fluctuation with a mixture of joint activation and joint silencing. The remaining pairs (34/316, 10.76%) exhibited preferential upward-biased direction, tending to become active together. Differences in directional preference did not predict the MI score (Mann-Whitney, U=3976, p=0.111, 281 bi-directional pairs *vs.* 34 upward-biased pairs). The occurrence of co-fluctuating L6b^Ctgf^ pairs likely reflects shared inputs that can promote either joint activation or joint inhibition (echoing our findings from *ex vivo* electrophysiology; *vide supra*, **Fig.S3**).

In brief, in awake freely moving mice, L6b^Ctgf^ neurons operate not through recurrent or oscillatory dynamics, but via flexible activity patterns maintained by a shifting subset of active cells. This dynamic baseline state provides a reference for interpreting population activity changes during more structured behaviors.

### L6b^Ctgf^ neuronal activity is modulated by the external environment and time-locked to behavior

Having defined the spontaneous operating mode of L6b^Ctgf^ neurons, we examined how their activity changes when the animal engages in behaviors that naturally recruit the barrel cortex. During active tactile exploration, whisking patterns are adjusted to enhance the acquisition of information^63^. Correspondingly, barrel cortex neurons exhibit characteristic activity signatures that reflect the active sensing process^64^. Recent work further shows that the SSp-bfd activity also reflects dimensions of whole-body positioning, including head-body angle and locomotor state^65^. By comparing distinct but complementary behavioral actions that emphasize each modality, we were able to dissociate the contributions of individual sensorimotor modalities and evaluate how each one shapes local L6b^Ctgf^ activity.

First, we analyzed calcium dynamics during rearing, a behavior used for information gathering^66^. When upright, mice whisk, but their whiskers remain in open air and therefore no tactile feedback is expected^67^. We annotated 95 rearing occurrences (only cases of unsupported rearing with full body extension; 24±3/mouse, 191 cells). For each neuron, we computed the average activity within a 10s window containing the moment of maximal upright extension. At the population level we observed no time-locked changes in L6b^Ctgf^ activity (**Fig.6C**). To determine whether specific neurons showed rearing-related modulation, we compared each neuron’s activity during rearing to its pre-bout baseline and assigned it to one of three response classes: *non-responsive, activated,* or *inhibited*. During rearing, response classes were represented in comparable proportions (non-responsive: 36.6%, activated: 33.0%, inhibited: 30.4%; **Fig.7A**, pie chart). Among the activated subset, the z-score signal peaked within the first 1.7s following maximal rearing height (z-score 55.45±9.01; **Fig.7A**, orange line), while inhibited neurons reached a trough and remained highly suppressed throughout that same period (z-score-33.54±3.49; **Fig.7A**, blue line), beginning to recover approximately 2.3s after maximal rearing height. Rearing events lasted 3.06±0.38s, beginning 0.80s before (front limbs disconnect from floor) and terminating 2.19 to 2.32s after maximal vertical extension. Therefore, activated neurons peak during the descent phase of rearing, and inhibited neurons begin to recover only after rearing terminates, *i.e.,* all four limbs re-establish floor contact.

**Figure 7.**
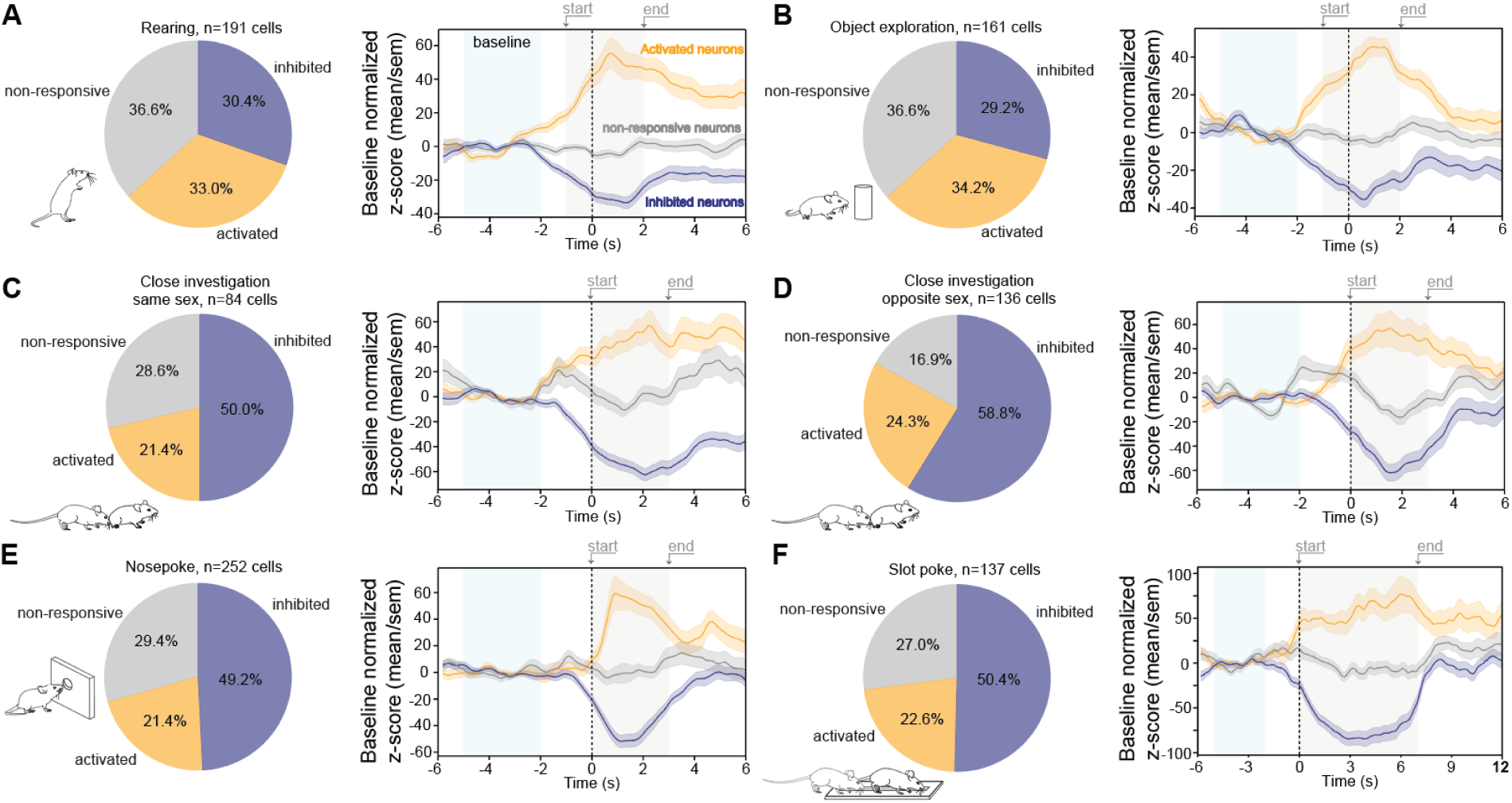
Behavior-dependent increase in the proportion of inhibited L6b^Ctgf^ neurons in the SSp-bfd. (A-F) Calcium dynamics. Left, proportion of cells classified as inhibited (blue), activated (orange) or non-responsive (gray). Right, behavior aligned average peri-stimulus time histograms for each cell class. Period used for baseline z-score correction (light blue shade) and window of time used for response class identification (light gray shade). Windows adjusted to behavioral bout duration. Lines represent the baseline normalized class response, mean±SEM z-score. Four mice. Same animals in all behavioral tests. SSp-bfd, primary somatosensory cortex barrel field. See also **Figure 6** for overall population activity heatmaps and **Figure S5**.

In contrast to rearing, during object exploration the animal maintains a grounded posture and actively protracts its whiskers forward, with object contact typically occurring at distances below 2cm from the snout^68^. Accordingly, we annotated only events in which the mouse oriented its head toward the object and approached (nostril tip) to within 0.8cm of the object surface (42 events, 10±2/mouse, 161 cells). We did not impose a strict definition of behavioral onset or offset; instead, we annotated the moment of maximal proximity to the object and analyzed the 10s surrounding window. Once again, population activity did not exhibit time-locked modulation (**Fig.6D**), and response classes were equally distributed (non-responsive: 36.6%, activated: 34.2%, inhibited: 29.2%). Both activated and inhibited subsets reached their respective peak and trough values within the same half-second window (activated: peak at +2.0s, z-score 45.49±5.16. Inhibited: trough at +1.6s, z-score-35.56±4.25)(**Fig.7B**).

Next, we examined L6b^Ctgf^ activity during social investigation. The social target provides strong olfactory cues that can modulate respiratory frequency and, in turn, influence whisking dynamics, creating a richer multisensory context known to recruit additional neurons in the barrel field^69^. We assessed activity during face-to-face interactions, which were comparable in posture to object exploration (24 events, 6/mouse, 172 cells). Face-to-face interactions produced no overall change in population activity (**Fig.S5B**), nor in the distribution of response classes (non-responsive: 41.9%, activated: 32.0%, inhibited: 26.2%). We noted that the response magnitude for active (peak at 0.7s, z-score 64.33±9.13) and inhibited (trough at +1.9s, z-score-54.83±6.39) neuronal classes was larger during face-to-face interactions during novel object exploration (z-score amplitude; ±40 object exploration *vs.* ±60 face-to-face interaction; **Fig.7B** *vs.* **Fig.S5C**). Next, we evaluated investigatory bouts in which the subject explored the conspecific’s anogenital area, adopting a characteristic posture with the head positioned beneath the tail of the stimulus animal. We separately evaluated anogenital sniffing toward same-sex and opposite-sex conspecifics (opposite sex; 24 events, 10±4/mouse, 136 cells. Same sex; 22 events, 9±3/mouse, 84 cells). The duration of physical contact between the nostril and anogenital area was brief (opposite sex 0.83±0.25s; same sex 0.62±0.24s). Yet, during anogenital sniffing, we observed a sharp decrease in overall population activity (**Fig.6E**). Response classifications shifted toward the inhibited class, which now comprised at least half of all neurons (50.0% opposite sex and 58.8% same sex). The average z-score of the inhibited class had large amplitude changes (z-score-62.60±4.98 and-61.92±7.03, respectively), with troughs at +2.1s for same-sex and 1.5s for opposite-sex. The activated subset aligned in amplitude (z-score 56.97±12.17 and 57.03±14.42) and timing (peak at 2.3s and 1.5s). Both neuron classes started shifting from baseline approximately 1s before contact was established, and started drifting back to baseline 1s after contact was lost (**Fig.7C-D**). We interpreted these changes as being aligned with the action of placing the head beneath the tail of the stimulus mouse.

We hypothesized a similar silencing could occur in conditions that involve whisker deflection but lack a social component. For this analysis mice explored a chamber containing a nosepoke-port, and we analyzed bouts in which the mice fully inserted their nostril into the port (93 events, 23±5/mouse, 252 neurons, nostril max depth 0.8±0.1cm, bout duration of 2.4s entry to exit). Again, we detected an attenuation in L6b^Ctgf^ population activity (**Fig.6F**) with half of the neurons classified as inhibited (trough occurring 1.2s after entry-start, z-score of-51.83±4.64; **Fig.7E**). Notably, the inhibited class maintained a flat minimum for the duration that the nostril remained in the poke-port (**Fig.7E**, blue line). The active class was composed of a minority of neurons (21.4%), reaching peak z-score (59.65±12.47) at max nostril insertion depth (peak at 0.9s).

To test whether L6b^Ctgf^ silencing is indeed maintained for the duration of nostril slot insertion, we sought to prolong the behavior by allowing animals to explore an elongated and narrow slot (slot poke; 22 events, 5-6/mouse, 137 neurons). Slot exploration took on average 4s (**Fig.S5D**) and was indeed accompanied by a longer period of L6b^Ctgf^ population silencing (**Fig.6G**), aligning with the longer behavioral engagement, and a stronger overall amplitude drop. Half of the neurons were classified as inhibited, reaching a minimum z-score of-84.79±8.30 with a trough at 3.9s. In contrast, the active subset comprised only 22.6% of neurons, with a peak z-score of 76.73±20.84 at 6s (**Fig.7F**). These observations indicate that slot sniffing is associated with both extended and enhanced silencing.

To conciliate data between behaviors, we compared effect sizes for all conditions. The normalized inhibited pool area was defined as the time-averaged magnitude of negative deflection in the z-score trace during behavior. Using this metric, we confirmed that slot sniffing exhibits the largest inhibition amplitude (normalized pool area, nAUC 69.8). Approximately 1.3x larger than what is observed during anogenital sniffing (opposite sex, nAUC 54.5; same sex nAUC 49.6) and 1.7x larger than nosepoke exploration (nAUC 40.9). Hence, L6b^Ctgf^ population silencing closely follows behavioral engagement, such that prolonging the action extends the duration of inhibition, and response magnitude scales with duration.

Our data support a model in which L6b^Ctgf^ silencing is associated with a whisker retraction-biased state that emerges when environmental features constrain whisker motion. During preparatory locomotor states, retraction of the whiskers also occurs and supports a functional link between whisker positioning and spatial attention^67,70^. For relating L6b^Ctgf^ activity and locomotor state, we used accelerometer data from a head-mounted inertial unit and identified transitions between immobility and locomotion (**Fig.S5H**). Transitions from immobility (when whisker retraction predominates) to locomotion (when protraction dominates followed by increased active whisking) were characterized by half of the neurons being inhibited although with small amplitude changes (nAUC 9.6; **Fig.S5I-J**). Overall, L6b^Ctgf^ neurons showed a higher probability of being active during sustained immobility, and decreased this probability during transitions from immobile to mobile states (**Fig.S5K**).

### Robust cross-behavior stability of inhibition as a L6b^Ctgf^ response mode

Given that all analyses were performed within the same cohort of animals, we were able to conduct a final assessment of response modality across behaviors. We selected the four behaviors in which an overall population inhibition occurred (anogenital sniff [opposite sex; same sex], nosepoke and slot poke). To evaluate persistence of cell response modality, class stability was compared to a random-transition null model based on overall class frequencies within these behaviors.

Non-responsive cells exhibited the greatest state variability, with a 0.71 probability of changing classification across behaviors, and activated cells showed a similar dynamic with a 0.66 probability of response modality change. Relative to their expected null stabilities (0.35 for non-responsive and 0.25 for activated cells), both classes were less stable than predicted by chance. In contrast, inhibited cells showed the strongest cross-behavior stability, with a probability of remaining in the same class of 0.62. This represents a +0.22 increase above expectation (null 0.40), showing that cells which are inhibited in one behavior are also inhibited in another more than 60% of the time (**Fig.S5E-G**). These results demonstrate that inhibited class cells form a robust and behavior-invariant population, whereas activated and non-responsive cells display intrinsically dynamic and context-sensitive response profiles.

## DISCUSSION

Here, we unveil several new aspects of a large subpopulation of L6b PSNs, providing convergent evidence that these neurons are *bona fide* members of the postnatal cortical network with unique specialized features and functions. We confirm the utility of CTGF as a selective and sensitive genetic identifier of L6b neurons. We show that L6b^Ctgf^ neurons occupy a continuum of electrical identities, and demonstrate salient connectivity patterns within the local cortical circuit as well as across regions. In awake freely moving mice, we identify a surprising relationship between somatosensory L6b^Ctgf^ neuronal activity and exploration of the environment.

The recognition of molecular diversity among PSNs^47,71^ has been vital to the last decade’s progress in characterizing these neurons as active members of the cortical circuit. *Ctgf* offers an attractive molecular handle in this work due to its restricted expression to L6b in the brain^40,47,72^. Other commonly used L6b markers, such as *Drd1a*^28,39,73,74^, are more problematic in this regard as their expression spans multiple cortical layers precluding clear laminar delineation^75,76^. Of note, Ctgf also delineates L6b in primates^41,77^, suggesting its utility for cross-species comparison. We confirm earlier reports supporting the utility of Ctgf-Cre mice for L6b studies^28,29,39,78^, and further demonstrate regional variations in cell density and layer thickness of this population across the cortex, with the highest density observed within the SSp. Our electrophysiological characterization of SSp-L6b^Ctgf^ neurons did not identify strict categories, but rather a continuum of properties, complementing previous findings from ‘non cell-type-specific’ studies with L6b neurons showing diverse firing properties^24^. Such diversity likely reflects the fact that *Ctgf* marks essentially all L6b PSN subclasses, albeit with incomplete penetrance within each class, as supported by recent transcriptomic data^43,79,80^. In addition, the heterogeneity in expression of the PSN markers including *Ctgf*, Neurexophilin 4 (*Nxph4),* and Complexin3 (*Cplx3)* appears to be further shaped by cortical modality with preferential enrichment of *Ctgf* in the somatosensory cortex, and *Drd1a* in the visual cortex.

Connectivity studies of L6b have shown substantial cortico-cortical projections^38,81^. Thus, with regard to input the largest innervation of L6b^Ctgf^ neurons, apart from local contributions, arrives from the corresponding contralateral cortex^28^. Our analysis, focusing on L6b^Ctgf^ output and taking advantage of a dual-color viral construct, allowed us to separate fibers of passage from axonal terminals. Using a brain-wide mapping strategy, we show that L6b^Ctgf^ neurons in SSp and MOp are characterized by origin-specific, ipsilaterally restricted projection patterns. Through distinct regional and laminar targeting, they form reciprocal modality-matching corticocortical connectivity. This arrangement resembles what has been described for cortico-cortical L5^82^ and L6a pyramidal neurons^83^, except L6b^Ctgf^ do not have callosal projections. This lack of interhemispheric efference has been observed with other PSN markers^78^.

Thalamic projections were also origin-site specific. For instance, L6b^Ctgf^ neurons residing in the MOp showed no thalamic projections, whereas those residing in the SSp projected to thalamic sensory motor-related areas. We did not find projections to the higher order thalamic nuclei from SSp-L6b^Ctgf^ neurons, setting Ctgf neurons apart from Drd1a layer 6 neurons, which innervate the posterior medial thalamus^38,39^. Further indication that L6b^Ctgf^ neurons maintain origin-site specificity comes from studies of the entorhinal-L6b^Ctgf^ population, which projects to the hippocampal CA3^29^. Interestingly, neither SSp-nor MOp-L6b^Ctgf^ fibers project to the hippocampal formation, though we did observe hippocampal arbors in our double transgenic tdT line which labels all *Ctgf* neurons.

The preferential presynaptic bouton distribution over some cortical layers raised the possibility that L6b^Ctgf^ neurons would connect selectively to specific cell types, while avoiding or making weak connections to others, a principle described for other cortical neurons^84,85^. However, this does not seem to be the case. Local optogenetic activation of L6b^Ctgf^ fibers in both SSp and MOp showed that they form functional monosynaptic connections with both pyramidal cells and interneurons, with high probability, especially in L2/3 and L5 where synaptic bouton density was high. (Notably, however, we found no direct synaptic L6b^Ctgf^-L6b^Ctgf^ connections, in agreement with Zolnik *et al*.^28^). Interestingly, our analysis of presynaptic strength shows that L6b^Ctgf^ neurons have short-term facilitating responses. This indicates that L6b^Ctgf^ neurons are more efficient in exciting target cortical neurons when spiking in rapid sequence. Additionally, we identify multispike activity intermingled with single spikes in half of all active L6b^Ctgf^ cells, suggesting that these neurons are capable of adapting their firing pattern to modulate the efficacy with which they entrain postsynaptic partners^86^. Indeed, our electrophysiological characterization reveals their glutamatergic nature, which is in line with labeling studies showing these neurons do not co-label with GABAergic markers^28,42,71,81^.

Previous studies of intracortical L6b connectivity have emphasized its targeting of pyramidal output neurons^39^ (see also^28^). However, we identify a substantial targeting of inhibitory interneurons across layers. Adding a downstream inhibitory component to the model can have significant consequences for the role of PSNs in cortical network dynamics. For instance, work on WM PSNs has shown their ability to recruit blanket inhibition which leads to desynchronization of cortical network activity^36^. This latter mechanism has been evoked as a means for these cells to shift the cortical circuit from a slow-wave sleep-like state to a pattern typical of sensory processing in the awake state^33^. With the circuit arrangement of PSNs clarifying, questions can now be asked about the role of this “layer with no known function”^31^. Important insight has been offered by recent optogenetic studies that suggest a role in increasing wake-like activity^39,73^. However, exogenous activation, while highly valuable, does not necessarily reflect physiological activity patterns. It is therefore important to also determine the dynamics of a specific population as aligned to behavior. To date, the few *in vivo* recordings of PSNs available have been restricted to head-fixed animals and without genetic specificity^25,87^.

Here, we characterize the activity of genetically defined PSNs in freely moving animals exploring their environment. *In vivo* calcium imaging of SSp-L6b^Ctgf^ barrel field neurons revealed that these cells are typically active during spontaneous behavior; however, population silencing emerged as the predominant response pattern during behavioral engagement. Under conditions expected to progressively recruit SSp-barrel circuitry, this silencing was tightly time-locked to behavioral onset and scaled with behavioral duration. Taking the conjoint of our findings and PSN literature a model now emerges where L6b^Ctgf^ PSNs, through their extensive intracortical connectivity with both pyramidal neurons and inhibitory interneurons, and previously demonstrated capacity for desynchronizing the cortical network^39^, could prime the cortical area for engagement once a relevant stimulus arrives, as a mechanism for “alertness”. Once activated by a salient stimulus of matched modality the majority of these neurons disengage, limiting access to the local cortical network so that it can process the salient signal. This would guarantee filtering out subsalient information.

Sparsity in activity of the SSp-bfd has been shown with less than 10% of excitatory neurons firing in response to whisker deflection, an effect considered to be mediated largely by a fast-spiking subtype of GABAergic inhibitory neurons^61^. In our data, the persistence of inhibited class neurons across several behaviors indicates that being silenced represents a true response mode of the L6b^Ctgf^ population. Approximately half of L6b^Ctgf^ neurons receive consistent behavioral-coupled inhibition when exploratory actions involve whisker deflection. By suppressing a large and consistent fraction of this population, this process may enhance contrast between salient information and the competing background signals. Ultimately suppressing alternative cortical dynamics^88^ which would otherwise be facilitated by the glutamatergic input of L6b neurons^73^.

While the population response followed behavior-linked silencing, a smaller subset of L6b^Ctgf^ neurons is activated with similarly locked temporal dynamics. Further with the model speculated above, this subset may provide excitatory drive to cortical neurons through the extensive cortico-cortical connections demonstrated here, facilitating encoding of relevant features via gain modulation^89^, whereas the projections to the primary thalamus would provide feedback information of cortical state activity^90,91^. Of note, the composition of the activated minority changes dynamically across behaviors. This flexibility in response to the paradigm applied could allow for a behavioral specific population representation. The likely candidates to coordinate selective L6b^Ctgf^ inhibition are interneurons residing in the local L6a network^28^.

The present study combines insights on the diversity, connectivity, and behavioral-linked activity patterns that cumulatively suggest a role for PSNs in modularly adjusting cortical state to optimize information gathering during exploration. These results suggest several exciting directions for future work, including addressing how the silenced and activated subsets of L6b^Ctgf^ neurons are dynamically selected in a behaviorally dependent manner, determining the response of L6b neurons to exogenously applied whisker deflection and exploration of L6b responses in other cortical regions as their respective modality is engaged.

## RESOURCE AVAILABILITY

### Lead Contact

Further information and requests for resources and reagents should be directed to, and will be fulfilled by, Christian Broberger.

### Materials Availability

This study did not generate new unique reagents.

### Data and Code Availability

Code used for analysis is available at [Zenodo: 10.5281/zenodo.19002257].

## Acknowledgments

This research was made possible by generous support from the Swedish Research Council (2021-00671_VR), internal funds from Stockholm University to C.B., and R.K. was supported by a Karolinska Institutet PhD fellowship. We thank the Imaging Facility at Stockholm University (IFSU) for assistance with confocal microscopy and the Experimental Core Facility (ECF) are acknowledged for their expert services. We acknowledge the Viral Vector Facility (VVF) of the Neuroscience Center Zurich. The authors thank all members of the Broberger laboratory for helpful discussions, and Dr. Paolo Medini for experimental suggestions. We express our gratitude to Drs. Jimena Ferraris and Paul Williams for management of animal colonies.

## Author Contributions

Project administration, Funding acquisition and Supervision, CB. Conceptualization, CB, RK, DM. Validation and Resource management, RK. Data curation, RK, DM, AL. Formal analysis and Visualization, RK, DM, AL, ON. Methodology CB, RK, DM, AL, ON. Investigation, RK, DM, AL, ON, JL. Software, DM, AL. Writing – original draft, CB, RK, DM, AL. Writing – review & editing, CB, RK, DM, AL.

## Competing Interests

The authors declare no competing or financial interests.

## Declaration of generative AI and AI-assisted technologies

During the preparation of this work, the authors used GitHub Copilot in order to curate and annotate scripts used during the analysis. After using this tool, the authors reviewed and edited the content as needed and take full responsibility for the content of the publication.

## STAR METHODS

Details of the design for each experiment, including within-and between-subject factors as well as full descriptions of variables required for independent replication are presented in each method’s subsection. Further information such as number of animals, genetic cross, sex, age, number of brain slices or cells evaluated per animal can be seen in figure legends and **Table S1** (groups and detailed stats), **Table S2** (inclusion/exclusion criteria), and **Table S3** (statistical summaries), **Table S4** (list of anatomical areas and acronyms).

## EXPERIMENTAL MODEL AND STUDY PARTICIPANT DETAILS

### Animals

The study was performed in heterozygous Ctgf*-*Cre mice (B6.Cg-*Ccn2^tm1.1(folA/cre)Hze^*/J, Jackson #028535)^92^. In these mice, Cre is degraded unless trimethoprim (TMP) is administered (Sando et al., 2013). To induce Cre recombinase activity, TMP was injected intraperitoneally (180-250µg/g, diluted in saline) after weaning (postnatal day 26-28). Ctgf-Cre mice were maintained heterozygous by backcrossing with C57BL/6J mice (Jackson #000664). For anatomical and electrophysiological experiments, Ctgf-Cre mice were crossbred with floxed-tdTomato mice (Gt(ROSA)26Sor*^tm^*^14^*^(CAG-tdTomato)Hze^*/J, Jackson #007914). The resulting double transgenic line tags neurons expressing *Ctgf* with the red fluorescent reporter protein, tdT.

Mice were group-housed (3-5/group) and kept on a 12h light-dark cycle (lights on at 7am) with food and water *ad libitum* (23-24°C, 45-65% humidity). Experiments were performed during the light phase of the cycle, and both sexes were used. Experiments were performed in accordance with the EU Directive 2010/63/EU, and approved by the local ethical committee.

## METHOD DETAILS

### In situ hybridization

Following the induction of deep anesthesia, Ctgf-Cre mice were sacrificed by decapitation, and the brains were rapidly dissected and frozen on dry ice. Coronal sections (14μm thick) were cut on a cryostat (Cryostar NX70, Thermo Scientific) and mounted onto SuperFrost glass slides (Epredia). Sections were fixed by immersion in 4% paraformaldehyde diluted in Phosphate Buffered Saline (PBS), and target mRNA was detected using the RNAScope® protocol (ACDBio). In brief, sections were incubated with *Ctgf* (314541-C1) and *Cre* (312281-C2) probes (1:50, C2:C1) for 2h at 40°C, followed by 30min incubation with signal amplifiers (40°C). *Ctgf* probes were labeled with Cy3 (Opal 570, 1:1500, Akoya Bioscience), and *Cre* probes were labeled with FITC (Opal 520, 1:1500, Akoya Bioscience). Twelve brain sections from six mice (two/mouse) were imaged by confocal microscopy (LSM800 Zeiss, 40X, Plan-Apochromat, 0.95NA) and co-expression analysis was performed using Fiji (v1.53q). To describe neuronal distance to the white matter, we segmented *Ctgf* expressing neurons and created a binary mask by thresholding with the triangle method^93^. Next, we extracted the x,y position of each neuron and measured the shortest distance between its centroid and the dorsal border of the white matter. For layer thickness we calculated a bivariate fit of x,y positions and extracted the residual sum (distance between centroids and the fitted curve) as a measure of neuronal dorsoventral spread. All sections were included in this analysis.

### Stereotaxic virus injections and lens implantation

Ctgf-Cre mice were anesthetized with isoflurane (3.5%) and placed in a stereotaxic frame (Stoelting). Following an incision to expose the skull and a unilateral craniotomy, a glass micropipette (PC-100 puller, Narishige), front-filled with the virus construct, was lowered into the brain. Viruses were slowly infused (200nL, 50nL/min).

The following viruses were used: for anatomical experiments AAV5-hSyn1-DIO-mEGFP-Syp1-mRuby (VVF-Zurich #V303, 6.7×10^12^vg/mL), or AAV5-pCAG-DIO-EGFP (Addgene #51502, 1.3×10^13^vg/mL), for optogenetic manipulation AAV5-EF1a-DIO-hChR2(H134R)-mCherry (Addgene #20297, 1×10^13^vg/mL) and for *in vivo* calcium imaging AAV-DJ-hSyn1-DIO-jGCaMP8m (VVF-Zurich #V628-Dj, 8.0×10^12^vg/mL).

Injection coordinates (mm, reference to Bregma): Primary somatosensory area (SSp; AP-0.59, ML-2.80, DV-2.4), primary motor area (MOp; AP 0.8, ML 1.35, DV-2.2). Experiments were initiated four weeks after surgery.

All material used for *in vivo* calcium imaging were acquired from Inscopix. Lens implantation was performed three weeks after viral injection. A head screw (Pinnacle #8209-0.10) was placed in the frontal bone to provide implant stability. The dura was removed, and a 0.5mm depth medial plane track (parallel to the virus injection site and perpendicular to the skull) was made using a straight-edged dissection knife (FineScience #10055-12). Next, a prism gradient index (GRIN) lens (1×4.3mm) was slowly inserted (0.05mm/min), with the triangular imaging surface facing towards the SSp (most caudal lens tip at AP-1.2, ML-2.0, with final DV position based on cell visualization). The lens was secured to the skull with adhesive primer (Optibond FL, Kerr), cement filling (Tetric Evoflow, Ivoclar) and covered with a plastic cap to protect its surface. Two weeks after implantation, the miniscope baseplate was mounted and the head plate painted in black. One week later, the field of view was inspected in the awake animal, and mice in which fluorescence variation could be observed with low LED irradiance (<0.8mW/mm^2^, 455nm) were selected for behavioral analysis. Implantation site was confirmed *post mortem*.

### Immunostaining and Anatomical analysis

Mice were deeply anesthetized with pentobarbital (200mg/kg) and transcardially perfused with cold PBS (4-8°C, 0.05M, Gibco) and heparin (10U/ml) followed by a 4% paraformaldehyde solution (pH 7.4, Histolab #HL96753). The brain was immersionfixed for 3h. The tissue was kept at-20°C in cryoprotectant solution until sectioning (v/v): ethylene glycol 30%, glycerol 30%, 0.2M phosphate buffer 30%, and distilled H_2_O 10% at pH 7.4. Brains were sectioned on a vibratome (Leica VT1000; coronal sections 60µm, sagittal sections 80µm). Sections were incubated for 1h at room temperature (RT) with a blocking solution of 0.5% Triton-X100 in PBS (PBS-T) and 5% donkey or goat serum (Invitrogen). Primary anti-DsRed (1:1000, Takara #632496) and anti-GFP (1:750, Aves Labs #GFP-1020) antisera were applied in solution (PBS-T, 1% serum) together with nuclear counterstaining (NeuroTrace 435/455, 1:500, Invitrogen #N21479) and incubated overnight, at RT. After washing (3×20min, PBS-T), fluorophore-coupled secondary antibodies (PBS-T, 1% serum), Alexa-594 anti-rabbit (1:500, Invitrogen #A21207) and Alexa-488 anti-chicken (1:500, Invitrogen #A11039) were incubated for 2h at RT. Sections were washed (2×15min, PBS) and mounted (ProLong Gold, Invitrogen #P36930). Tile imaging of full sections were taken with a fluorescence microscope (AxioObserver Z1, Zeiss, 20X, Plan-Apochromat, 0.8NA).

Both coronal and sagittal sections were registered to the Allen Brain Atlas CCFv3^53^ using the ABBA whole-brain mapping pipeline^94^. For each section, the anteroposterior (coronal slices) or mediolateral (sagittal slices) position was identified. Next, we used affine and spline registration approaches (NeuroTrace channel, affine; 40µm sampling, with Mattes mutual information algorithm for optimization, and spline; 20 reference points, re-sampling of 15µm) to achieve an optimal registration overlap. Segmentation of neuron arborization and synaptic boutons was performed using the QuPath bioimage software^95^. To segment neuronal arborization, we trained a multilayer artificial neural network with 1 392 annotated regions containing fibers and 398 regions devoid of fibers (extracted from all animals). The main parameters used for training were; gradient magnitude (detection of edges), structure tensor max eigenvalue and Hessian eigenvalue (detection of string shaped objects) and structure tensor coherence (detection of objects with preferential orientation). After training, the software could segment fibers based on the pattern of fluorescence and its intensity (98% accuracy). For detection of synaptic boutons, we implemented pixel segmentation based on shape, size, and local fluorescence level. Segmented pixel clusters were annotated as presynaptic boutons only when there was colocalization with a fiber. The resulting data set contained, for each brain region in each slice, the area covered by fibers and the number of synaptic boutons. Sagittal sections were used to count the number of *Ctgf*-expressing neurons across the brain, as well as for complementary visualization of axonal arbor rostro-caudal extent.

### Brain slice preparation for electrophysiology recordings

Mice were deeply anesthetized with isoflurane and decapitated. Brain slices were prepared using an adaptation of a previously reported procedure^26^. The brain was removed and placed in ice-cold (2-4°C) oxygenated (O_2_ 95%, CO_2_ 5%) slicing solution consisting of (in mM): sucrose (214), KCl (2.5), NaHCO_3_ (26), NaH_2_PO_4_ (1.3), D-glucose (10), CaCl_2_ (2), MgSO_4_ (2). Coronal slices (250µm) containing SSp and MOp were cut on a vibratome (Model 5100mz, Campden) and transferred to a holding chamber containing oxygenated artificial cerebrospinal fluid (aCSF, 35±2°C, pH 7.4). For characterization of electrophysiological properties, aCSF consisting of (in mM): NaCl (124), KCl (3.5), NaHCO_3_ (26), NaH_2_PO_4_ (1.25), D-glucose (10), CaCl_2_ (1), MgSO_4_ (1), was used. For optogenetics experiments aCSF consisting of (in mM): NaCl (127), KCl (2.0), NaHCO_3_ (26), NaH_2_PO_4_ (1.2), D-glucose (10), CaCl_2_ (2.4), MgCl_2_ (1.3) was used. Following a minimum of 30min incubation, slices were transferred to the recording chamber and continuously perfused with oxygenated aCSF.

### Electrophysiological recordings

Electrophysiological characterization of L6b^Ctgf^ neurons was performed in the perforated patch configuration^96,97^. Patch pipettes had a resistance of 4-7MΩ and were filled with an intracellular solution containing (in mM): K-gluconate (140), KCl (3), HEPES (10), EGTA (0.2), MgCl_2_ (2), adjusted to pH 7.3 with KOH, 290mOsm. Neurobiotin (0.20%, Vector) and amphotericin B (200µg/ml, Sigma) were added to the internal solution. Liquid junction potential was 15.4mV and compensated. Capacitance values were extracted (membrane test, Clampex). De-and hyperpolarizing current pulses were applied as one second steps from-180 to 120pA with 20pA increments and 12s inter-pulse interval. Input resistance was calculated in response to-120pA current injections. For a detailed analysis of firing frequency responses, five randomly selected neurons received smaller increments (5pA increments,-150 to 135pA). Inclusion/exclusion criteria details are described in **Table S2**. For additional electrophysiological experiments in a whole-cell (single or paired) patch clamp configuration, amphotericin B was omitted from the internal solution, and Na_2_ATP (2mM) was added (recordings were collected >5min after membrane rupture).

Optogenetic experiments were performed in whole-cell configuration. The internal solution for these experiments contained (in mM): K-gluconate (140), KCl (10), HEPES (10), EGTA (1), Na_2_ATP (2) adjusted to pH 7.3 with KOH, 290mOsm. Neurobiotin (0.20%) was added on the experimental day. Photostimulation was performed with either wide-field illumination (light source: CoolLed 470nm, 1mW) through the microscope objective (W Plan-Apochromat, 63X, 1.0NA, 317µm illumination field), or via an optic fiber (200µm, 0.22NA, ThorLabs) placed on the tissue surface coupled to a light source (CNI 447nm, 1mW). Light pulses (1-10ms) were controlled by a digital output signal (DigiData 1440A, Molecular Devices). To determine the frequency-dependent spiking fidelity of ChR2-expressing neurons (mCherry+) we recorded in current-clamp configuration (optogenetic parameters: 10ms pulse, 2s duration, 5-40Hz, 1mW). To functionally assess the connectivity of L6b^Ctgf^ ChR2-expressing neurons with other cortical neurons, cells not expressing mCherry, were recorded across different cortical layers in voltage-clamp mode and held at-70mV (optogenetic parameters: 10ms paired-pulses, 50ms inter-pulse-interval, 1mW). Pharmacological manipulation was performed by bath application of blockers of fast synaptic transmission (gabazine SR95531 [GBZ, 10µM], tetrodotoxin [TTX, 1µM], 4-aminopyridine [4AP, 100µM], cyanquixaline [CNQX, 10µM], and D-2-amino-5-phosphonopentanoate [AP5, 10µM]. All drugs were acquired from Tocris. In all experiments, the pipette capacitance was automatically compensated and access resistance (R_a_) was continuously monitored. Neurons in which R_a_ changed >20% were excluded from analysis. The signal was amplified and low-pass filtered (10kHz, MultiClamp 700B, Molecular Devices), and digitized (DigiData 1440A, Molecular Devices). Data were acquired with Clampex and analyzed with ClampFit (v11.2, Molecular Devices). For assessment of neuronal location within the cortical layers (SSp and MOp), images were taken at the end of each recording session (Fluorescence and Differential Interference Contrast; DIC) with three magnifications; 63X, 20X and 5X. The location was confirmed using Fiji (v1.53q) to extract neuronal depth (from pial surface) and distance to the WM, followed by a normalization factor (distance: pia to WM) and projection of cell position onto the Allen Brain Atlas (CCFv3).

### Electrophysiological analysis

In perforated patch recordings, data were extracted for both L6b^Ctgf^ (tdT-expressing) and neighboring tdT-negative (L6b^Others^) neurons. After a 15min stabilization, baseline activity was recorded (5min), and neurons that did not fire any action potential (AP) were classified as silent (all else, active). To unbiasedly identify multispike events, we plotted a histogram of all interspike intervals (time between two AP peaks) for each neuron. Next, histograms were normalized to plot the global average. The observed bimodal distribution contained a col (lowest point between histogram peaks) at 43.5ms. Based on this, we defined a multispike event as any AP followed by another AP within 43.5ms (MATLAB vR2021a, MathWorks). Using this definition, we estimated the proportion of APs that occurred within trains and characterized their properties, including the number of APs per train and firing frequency.

We assessed passive properties of L6b^Ctgf^ active and silent neurons and compared them to L6b^Others^. To characterize APs for all neurons (including the silent class), we applied a 120pA depolarizing current pulse and analyzed the first ten elicited APs. The following features were extracted: AP threshold (mV), peak amplitude (mV), peak-to-peak frequency (Hz), max rise slope (mV/ms), rheobase current (pA), afterhyperpolarization (AHP) amplitude (mV) and time to peak (ms). Beyond AP characteristics, we also assessed sag and rebound properties. The sag potential was quantified using a-120pA hyperpolarizing current pulse, measuring the lowest deviation from baseline. Whereas the rebound potential was evaluated using a-180pA current pulse, defined as the highest peak of the positive deflection from baseline.

Spontaneous postsynaptic potentials in whole-cell recordings were detected using MiniAnalysis 6.0 (Synaptosoft). To generate cross-correlograms we compute the time difference between each detected postsynaptic potential (PSP). The resulting values were binned (-10 to +10ms window) with a 1ms resolution and z-scored to the mean and standard deviation value within that window (MATLAB 2024, MathWorks).

In optogenetic experiments, we evaluated spike fidelity upon different stimulation frequencies by calculating the percentage of evoked APs over the total number of light pulses (ten sweeps per frequency/cell). To evaluate the amplitude of light evoked excitatory postsynaptic currents (EPSCs, pA), all consecutive sweeps were averaged. Paired pulse ratio (PPR) was analyzed by dividing peak2 over peak1 amplitude^98^.

### Post-hoc identification of neurobiotin filled neurons

After electrophysiological recordings, the slices were transferred and incubated overnight at 4°C in 4% PFA/0.16% picric acid. After fixation, the slices were washed in 0.1M PBS and incubated with FITC-Avidin (1:2500, Invitrogen) in 0.1M PBS containing 0.6% Triton X-100 and 1% bovine serum albumin for 72h at 4°C. After washing with PBS, the slices were mounted in a DABCO medium and imaged with a confocal microscope (DMi8, Leica, 20X/0.75NA and 40X/1.25NA with Leica-Type^G^ immersion liquid).

### Calcium Imaging Experiments

Experiments used 4-to 5-month-old mice (n=4; two males, two females) implanted with a prism lens over the right SSp-barrel field (bfd). Animals were handled daily for five days before the first session. Six recording sessions (s1 to s6) took place over six weeks.

Two open-field chambers were used: a square chamber (30×30 cm) and a rectangular chamber (30×15 cm). In the square chamber, mice were recorded from below (transparent floor), in the rectangular chamber from above. Videos were acquired at 25fps and 1280×1024 resolution (GigE acA1300-60gm Basler camera, LMVZ4411 1/1.8, 4.4–11 mm/F1.6 Kowa lens). Illumination was held constant (100lux).

Each session began with a 10min baseline period in which the mouse was alone, followed by a 5-10min phase in one of the following conditions: mouse remains alone (s1), object exploration (s2), social interaction with a same-or opposite-sex conspecific (s3-s4; counterbalanced), nosepoke exploration (s5), or slot poke exploration (s6). In s2, the object was a plastic cylinder (a 50ml Falcon tube; 3cm diameter, 5cm height; 5min exposure). Conspecifics in s3 and s4 were C57BL/6 mice of same age, which had no prior contact with the subject (5min social interaction). In s5, animals were placed in a rectangular chamber where a nosepoke wall was positioned at one end, containing two identical 1.5cm openings spaced 2.5cm apart, with their lower edges 2cm above the floor. The opposite end of the chamber contained a top wall forming a shallow shelter (7cm cover), encouraging the animal to alternate between the nosepoke wall and the sheltered end of the apparatus (10min exposure). In s6, animals were tested with a transparent plexiglass panel (20×7cm) containing a central narrow slot (15cm long, 1cm wide, 0.9cm deep), positioned on the floor at the center of the square open-field. During s6 the test duration was adjusted so that each animal produced a comparable number of slot-exploration bouts (5-10min, min-max range).

*The following behaviors were annotated* (BORIS v8.27.4) by a researcher without access to calcium imaging data: (s1-5) **Rearing** events, occurring when no objects or conspecifics were present, identified as moments in which the subject raised on its hindlimbs. The event time was assigned at the point of maximal vertical extension. (s2) **Object exploration** was scored when the mouse approached the object with its head oriented toward it and all limbs on the floor. Events were marked at the moment of maximal proximity (<1cm distance between object and nostril). Bouts in which the animal failed to reach this minimal proximity were excluded from further analysis. (s3-s4) **Close investigation of a conspecific** was annotated in two forms. (i) When the subject approaches the conspecific, head directed towards it, and both animals sniff each other face-to-face. Annotated as a timepoint marking the moment of maximal proximity (as in object exploration). (ii) When the subject established contact between its nostril and the anogenital area of the visitor; in this case, the subject’s head was positioned below the tail of the conspecific, as verified from the below-view camera. The event time starts in the first frame with observed contact, and bout duration spanned from initiation to loss of physical contact. (s5) **Nosepoke exploration**, defined as bouts in which the mouse inserts its nostril into the aperture and withdraws it fully, with the behavior spanning from initial entry to complete retraction. The event time point marks the moment of maximal entry (average nostril depth 0.8±0.1cm). (s6) **Slot exploration**, defined as a continuous bout in which the animal lowers its nostril into the slot, moves along its length, and exits at the opposite end. For analysis, only bouts in which the subject kept its nostril within the slot for the entire traversal were included. For all behaviors, the ‘event time’ denotes the frame used to align behavioral actions with the calcium imaging data, serving as the zero point of the peristimulus interval.

For each animal and in each experimental day, a single raw calcium imaging video (10Hz) was recorded (Inscopix nVista, DAQ v2.3.0). Acquisition parameters were maintained across experimental sessions and an active commutator was used to prevent cable entanglement. The resulting calcium imaging movies were processed using the Inscopix Data Processing Software (v1.9.5.3874). First, defective pixels were fixed (3×3 median filter), and videos were spatially down sampled (factor 2, interpolation). Next, spatial bandpass filtering removed low and high spatial frequencies (0.5 and 0.005, respectively), and videos were motion-corrected. The baseline fluorescence value for each pixel, in each experimental day, was calculated as the average across all time frames of that video. Thereafter, regions of interest (ROI) were identified by Principal Component Analysis and Independent Component Analysis. Parameters as follows; average ROI diameter 13pix, block size 1000, convergence threshold 1e-05, max iterations 100. Detected ROIs were pre-accepted based on the number of components (n=1), a minimum ROI size of seven pixels, and signal-to-noise ratio (SNR) >3.0. Traces were denoised (first order autoregressive model, lag 5, noise range 0.25-0.5, spike SNR>3.5)^99^. At this stage, ROIs with roundness <0.2, were excluded to avoid instances of neuropil detection. After curation, approximately 17% of the initially detected ROIs were excluded from the data set. All remaining ROIs (n=255) were tagged as accepted neurons. To generate a session-spanning cell list for each animal, we performed longitudinal registration with a minimum correlation of 0.5. This criterion allowed us to match neurons across days and identify when the same cell was present in successive sessions. Recording sessions lasted 15-25 minutes, yielding a total of approximately 2 to 2.25 hours of data per animal.

*To identify neurons exhibiting time-structured calcium dynamics* (done in s1), each fluorescence calcium trace (ΔF/F) was first detrended using a 4^th^ order polynomial^100^. The normalized autocorrelation function was then computed for each trace. The most prominent peak (at lag>0) was extracted as the empirical measure of the neuron’s autocorrelation amplitude, and its corresponding recurrence frequency was calculated. To determine whether neurons displayed higher-than-chance autocorrelation, we generated a neuron-specific null distribution of autocorrelation peaks using a block-bootstrap procedure. Detrended traces were segmented into non-overlapping 5s blocks, and block order was randomly permuted 10^3^ times, preserving short-timescale correlations while disrupting long-range temporal structure. For each bootstrap surrogate, the analysis was repeated (as above), yielding a distribution of chance-level autocorrelation amplitudes. A neuron was classified as exhibiting significant recurring activity if its empirical autocorrelation peak exceeded the 95^th^ percentile of its bootstrap-derived null distribution and passed a Benjamini-Hochberg FDR correction (q<0.01) applied to all neurons recorded in the same animal.

*For the analysis of neuronal activity episodes* (done in s1), calcium fluorescence traces (ΔF/F) were high-pass filtered (2^nd^ order Butterworth, 0.01Hz) to remove any residual baseline drift. For each neuron, an activity threshold was defined directly from the filtered trace (F) using a robust median absolute deviation (MAD) criterion. The threshold (ε) was defined as ε = median(F) + 2 × MAD(F). Frames in which the fluorescence signal exceeded this threshold (F>ε) were classified as active. Contiguous active frames lasting ≥300ms were grouped and labeled as activity episodes. For each neuron, we quantified both the number and the duration of these episodes. To quantify population dynamics, we computed the instantaneous fraction of neurons engaged in an active episode using a 500ms sliding window. For each animal, the fraction of active cells was calculated for every window and then averaged to obtain the mean population activity fraction. To capture the strongest recruitment, we identified the peak window value for each animal, representing the maximum fraction of cells simultaneously active. Finally, we summed all time windows during which more than 70% of the population was labelled as active to provide a metric of cumulative duration at high-population engagement.

*For the analysis of neuronal co-fluctuating pairs* (done in s1), calcium traces were filtered as above. Frames with |F| < ε were set to zero (*i.e.,* non-fluctuating). The remaining values were converted into discrete non-zero signs (+1 for upward steps and-1 for downward steps). Two neurons were considered co-fluctuating only when both respected the following criterion: (i) both exhibited simultaneous sign episodes (+1 or-1, lasting ≥300ms) with overlapping segments of at least 200ms, and (ii) the total co-fluctuating time exceeded 1% of the recording duration. Pairs failing these criteria were excluded to remove brief, spurious co-occurrences before statistical evaluation. For each pair that met the criteria, we quantified the proportion of valid frames in which the two neurons fluctuated in the same direction (+1/+1 or-1/-1) or in opposite directions (+1/-1). These pairs were then compared against a surrogate null model generated by independently circularly shifting each neuron’s sign sequence (pair based, circular shift control, 10^3^ iterations). This procedure preserves each neuron’s temporal structure while removing genuine cross-neuronal alignment. For every surrogate, the same-and opposite-direction fractions were recomputed, yielding null distributions for both metrics. Observed values were converted to z-scores using the surrogate mean and variance, and statistical significance was assessed using the Benjamini-Hochberg FDR (q<0.01). Pairs passing this chance-level statistical evaluation were considered a *de facto* co-fluctuating pair (*i.e.,* temporally aligned above chance). The primary reported metric is the proportion of co-fluctuating pairs, relative to all theoretically possible pairs (within an animal). To estimate the expected number of detectable co-fluctuating pairs under a scenario in which all cross-neuronal temporal alignment is altered, we applied an additional circular-shift procedure at the population level. In this null control, we independently shifted each neuron’s sign sequence (population based, circular shift control, 10^3^ iterations), and the entire population was re-evaluated for co-fluctuating pairs. For every surrogate population, the full set of neuron-neuron pairs was tested using the same criteria. We provide a count of how many pairs would be deemed significant in a fully time-scrambled dataset (mean, 95%CI).

*To characterize whether co-fluctuating pairs showed a preferred directionality*, we quantified the relative proportion of time the pair spent locked in upward (+1/+1), downward (-1/-1), or opposite (+1/-1) sign co-fluctuation. For each pair, the relative contributions of these event categories were compared. If none of the three co-fluctuation categories accounted for the largest proportion, the pair was labeled bi-directional (reflecting coordinated timing despite variable sign relationships). Last, we computed mutual information (MI) to assess the extent of shared information between co-fluctuating pairs, capturing how knowledge of one neuron’s state constrains the possible states of the other (MI=1 indicates complete predictability; MI=0 indicates no predictive information).

*To quantify overall population activity during a behavior*, we first computed each cell’s mean activity from 5s before to 5s after each behavioral reference timepoint (note that for s6 the window was extended +5s). We then averaged these traces across all cells and plotted the overall population activity as a peri-stimulus line graph (grand average, mean±SEM). For single cell activity visualization, three options of min-max normalized heatmaps are shown. The first is sorted by overall activity level, placing the most active cells at the top. The second is sorted by the timing of each cell’s activity minimum, placing cells with the earliest trough at the top to reveal temporal structure in signal decreases. The third is sorted by the timing of each cell’s activity peak, highlighting temporal structure in signal increases. Because sorting enforces a monotonic ordering of event times, randomly timed cellular activity yields a diagonal band; whereas deviations from this pattern (*e.g.,* clustering or curvature) indicate structured temporal dynamics.

*To characterize how individual neurons responded during each behavior*, neuronal activity was classified into three response categories: non-responsive, activated, or inhibited. Classification was based on quantified changes in each neuron’s calcium signal during the behavioral bout relative to its immediate pre-behavior baseline. Baseline activity was computed over a fixed window extending from-5s to-2s prior to behavior onset. For each neuron, the mean fluorescence during the behavior window was compared to the baseline mean and standard deviation. Neurons were classified as activated or inhibited if the behavior-epoch mean exceeded or fell below the baseline mean by more than two baseline standard deviations, respectively; all others were classified as non-responsive. The implementation was dynamic, as the duration of each behavior window was defined by the actual behavior bout length. From this classification we report: (i) *class proportions*: for each behavior, we quantified the distribution of neuronal response types by calculating the proportion of cells classified as activated-during, inhibited-during, or non-responsive (proportion shown with pie charts). (ii) *Class response profiles*: to visualize the temporal dynamics of each response class, we computed activity as a peri-stimulus line graph (mean±SEM). (iii) *Peak and trough timing*: to compare response timing across classes, we extracted the time of the maximum (peak) or minimum (trough) signal within the behavior window. (iv) *Modality specificity across behaviors*: to evaluate whether neuronal response patterns were behavior-specific, we compared each neuron’s classification across behaviors. This allowed us to determine whether a neuron maintained a consistent response type (detailed on next paragraph).

*To quantify how neuronal response types reorganized across behaviors*, we constructed a cell-by-behavior classification matrix in which each entry contained the categorical response outcome (activated, inhibited, or non-responsive). Pairwise transitions were evaluated for all behavior combinations. For each pair, we tabulated the number of neurons transitioning from each class in behavior A to each class in behavior B, forming a 3×3 contingency matrix that was row-normalized to generate transition-probability matrices. Statistical dependence between classification distributions was evaluated using chi-square tests. A global transition matrix was then computed by pooling transitions across all behavior pairs. From this matrix, we derived class-specific stability probabilities (P_stay_, the probability of remaining in the same response class across behaviors). To determine whether neurons preserved their response class more often than expected by chance, we compared the empirical stability probabilities to a population-level null model. The overall population fractions of each response class served as the expected null probabilities under a model in which class assignments across behaviors occur independently of prior classifications. Empirical stability corresponded to the diagonal entries of the global transition matrix, and the difference between observed and expected values quantified class persistence beyond chance expectations. All stability metrics, expected-null values, and observed-versus-expected comparisons are reported.

*For locomotor state detection* the accelerometer data from the head-mounted inertial measurement unit (Inscopix, IMU) was used. To quantify movement, body acceleration (BA) was computed as the rectified sum of acceleration across all three orthogonal accelerometer axes, which eliminates dependence on sensor orientation relative to the animal’s reference frame. The resulting BA signal was low-pass filtered at 5Hz using a 3^rd^ order zero phase Butterworth filter and locomotor states were identified using a threshold-based detection approach adapted from prior studies^101,102^. In short, for each dataset, movement threshold was determined using Otsu’s method applied to the distribution of log-transformed BA values. This approach is effective because the log(BA) distribution is bimodal, reflecting periods of immobility and movement. Time points exceeding the threshold were classified as movement, whereas values below the threshold were classified as immobility. Locomotor onsets and offsets were defined by threshold crossings in the BA signal using fixed-duration state criteria. Transitions from immobility to movement required at least 2s of continuous immobility followed by at least 2s of continuous movement, whereas transitions from movement to immobility required the inverse sequence. The boundary between the two states was defined as the transition time point and used for temporal alignment in peri-transition analyses. Periods lasting longer than 4s of continuous immobility were used as a comparison time window. Locomotor transitions were analyzed from the first 10min of each behavioral session, during which the animal was alone in the square open-field chamber.

## QUANTIFICATION AND STATISTICAL ANALYSIS

Statistical analyses were conducted using Prism GraphPad (v9.3.1) or JMP analysis software (v14, SAS Institute). Normality was evaluated using D‘Agostino-Pearson test. For normally distributed data, t-test or one-way ANOVA followed by Tukeýs *post hoc* was used. For non-parametric data with two groups, Mann-Whitney. Otherwise, Kruskal-Wallis and Friedman tests followed by Dunńs *post hoc* were used. When specified, Benjamini-Hochberg FDR correction (q<0.01) was used. All statistical tests were two-tailed. Statistical significance is reported as exact values and, in the figures, as follows: *p<0.05, **p<0.01, ***p<0.001, ****p<0.0001. Values are reported in text as mean±SEM if not stated otherwise. Graphical data are presented as mean±SEM or mean±95%CI as indicated in the figure/legends.

Statistical details are reported in **Table S3**.

## SUPPLEMENTARY FIGURES

**Supplementary Figure 1.**
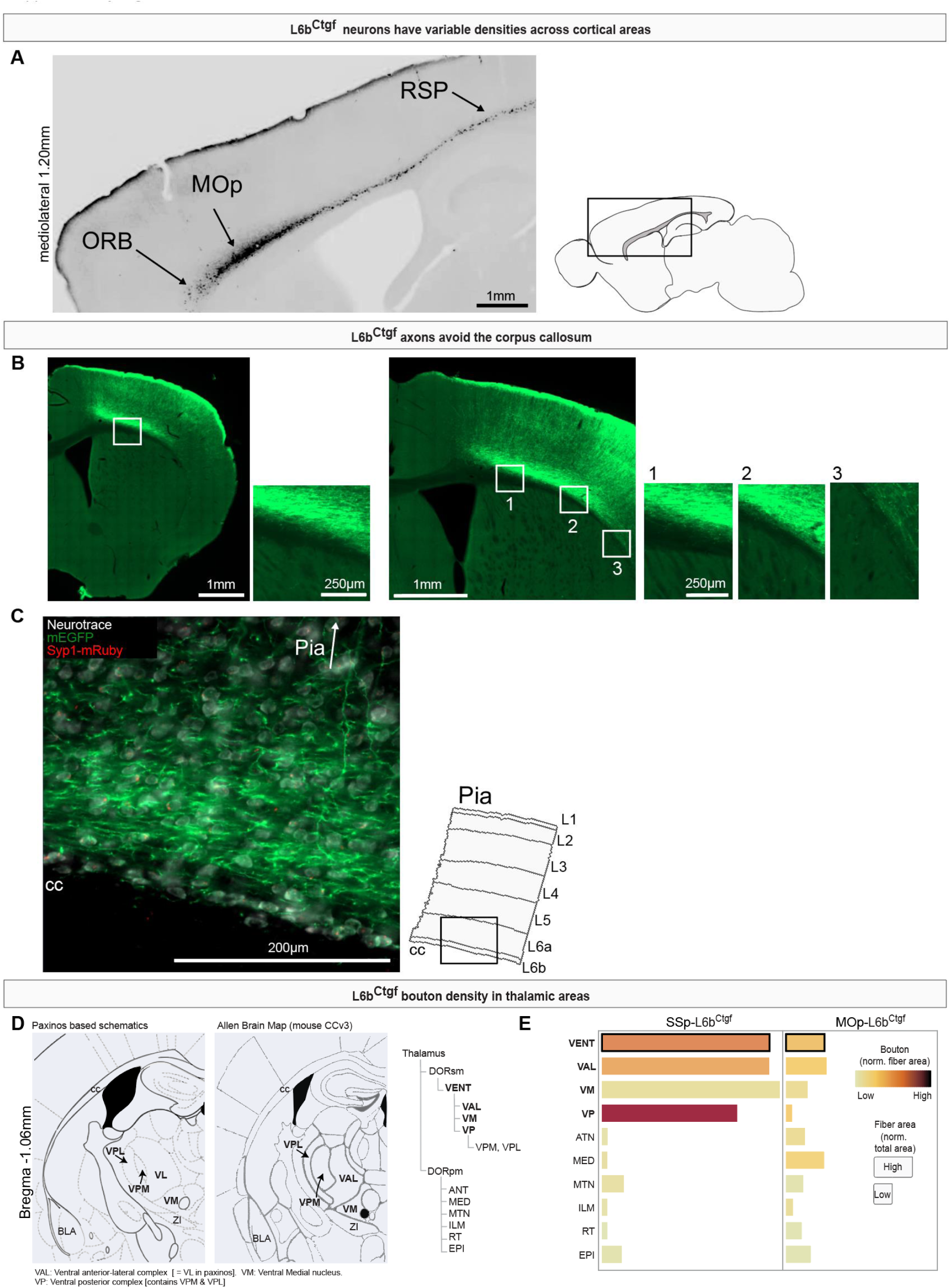
Connectome of L6b Ctgf-expressing neurons. **(A)** L6b^Ctgf^ neurons form a narrow band immediately above the white matter, with rostrocaudal variations in cell density. Left, fluorescence micrograph of a sagittal section with tdT-labeled cell bodies segmented (black; inverted grayscale). Right, schematic indicating the anatomical location of the section. **(B)** L6b^Ctgf^ axons avoid entry into the corpus callosum (cc). Photomicrographs show L6b^Ctgf^ neurons expressing EGFP (green) in SSp-L6b. Insets, higher magnification views in which the corpus callosum (cc) appears as a darker region devoid of EGFP signal. **(C)** Close up image of the SSp showing neurons (white, neurotrace), *Ctgf* fibers (green) and synaptic buttons (red dots). Right, schematic indicating the anatomical location of the imaged region. **(D)** Thalamic anatomical divisions and nomenclature. Left, divisions based on the Paxinos mouse brain atlas; right, corresponding divisions extracted from the Allen Mouse Brain Atlas, with the hierarchical organization of thalamic subregions shown. Note that the ventrolateral thalamic nucleus (VL) in Paxinos corresponds to the ventral anterior lateral complex (VAL) in the Allen atlas. **(E)** Quantification of thalamic innervation by L6b^Ctgf^ neurons. For each thalamic region, fiber density (box size) and bouton density (color scale) are normalized to total regional area. Left, mice injected in SSp; right, mice injected in the MOp. Data were processed as in **Figure 3F** showing pattern for cortical layers. **The following Acronyms from Paxinos are used in** (**D**, left): BLA, Basolateral amygdaloid nucleus. cc, Corpus callosum. VL, ventrolateral thalamic nucleus. VM, ventromedial thalamic nucleus. VPL, ventral posterolateral thalamic. VPM, ventral posteromedial thalamic. ZI, zona incerta. **All other acronyms follow the Allen Atlas nomenclature**: ATN, anterior group of the dorsal thalamus. DORpm, thalamus polymodal association cortex related. DORsm, thalamus sensory-motor cortex related. EPI, epithalamus. ILM, intralaminar nuclei of the dorsal thalamus. MED, medial group of the dorsal thalamus. MOp, primary motor area. MTN, midline group of the dorsal thalamus. ORB, orbital area. RSP, retrosplenial area. RT, reticular nucleus of the thalamus. SSp, primary somatosensory area. VAL, ventral anterior lateral complex of the thalamus. VENT, ventral group of the thalamus. VM, ventral medial nucleus of the thalamus. VP, ventral posterior complex of the thalamus. VPL, ventral posterolateral nucleus of the thalamus. VPM, ventral posteromedial nucleus of the thalamus. Related to **Figure 2** and **Figure 3**.

**Supplementary Figure 2.**
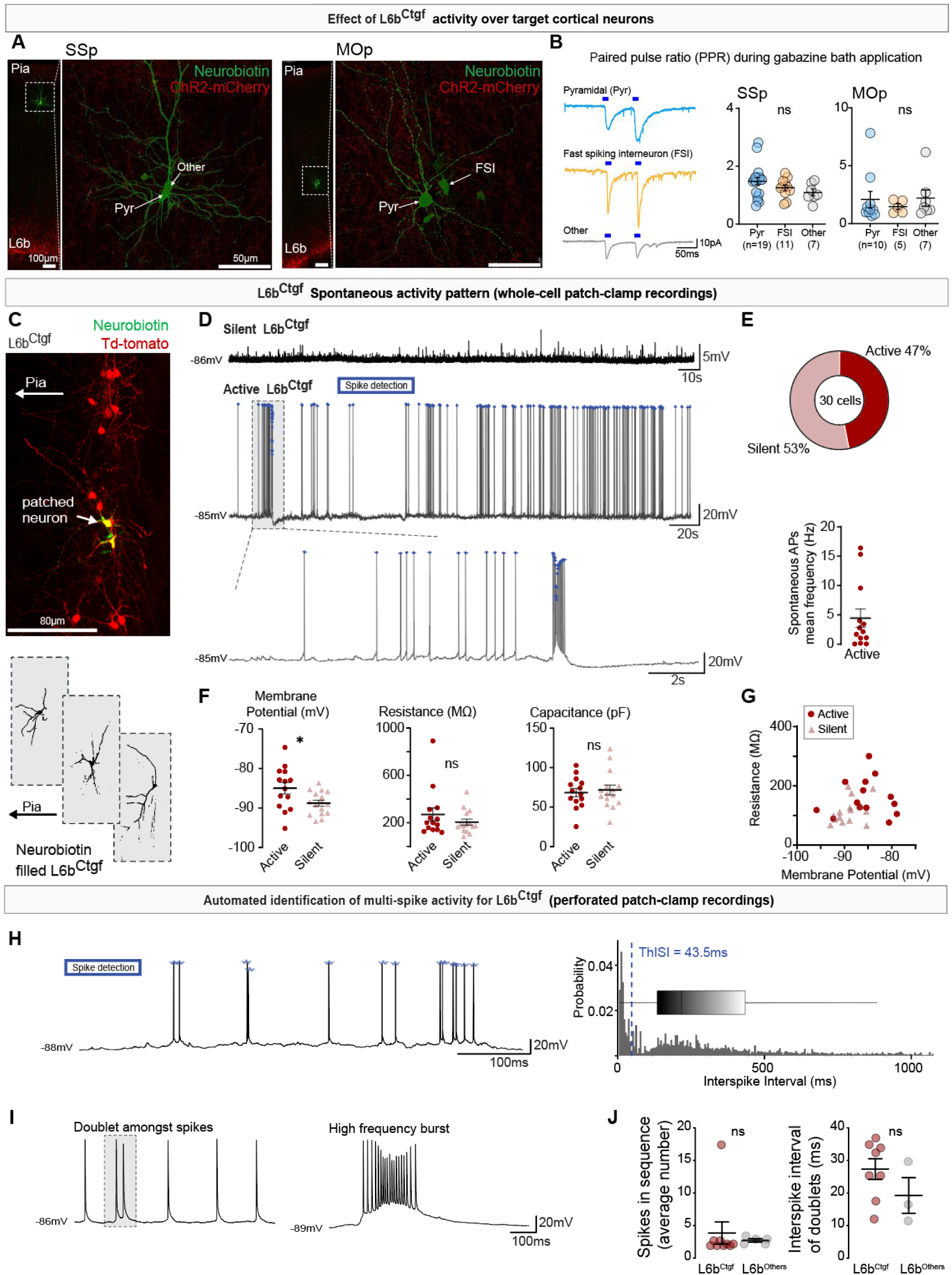
Paired-Pulse facilitation and heterogeneous intrinsic electrophysiological properties of L6b^Ctgf^ neurons. **(A)** ChR2 infected L6b^Ctgf^ neurons (red) project to, and form synaptic connections with other local cortical neurons. Representative confocal images show neurobiotin filled patched neurons (green). Patched neurons were recorded in acute ex-vivo cortical slices and responded to optogenetic activation of L6b^Ctgf^ terminals (Pyr, pyramidal. FSI, fast-spiking interneurons. ‘Other’, other interneurons). **(B) Paired-pulse facilitation of L6b^Ctgf^ outputs.** Optogenetically evoked EPSCs were elicited by paired photostimulation of L6b^Ctgf^ terminals with a 50ms inter-stimulation interval. The average PPR was consistently >1, indicating facilitating synaptic transmission from L6b^Ctgf^ neurons onto cortical targets in both SSp and MOp. Stats for SSp: Kruskal-Wallis H(2)=4.94, p=0.085. Mean±SEM (ratio) Pyr 1.48±0.13, FSI 1.26±0.1, Other 1.10±0.12. Stats for MOp: Kruskal-Wallis H(2)=0.47, p=0.80. Mean±SEM (ratio) Pyr 2.09±0.71, FSI 1.47±0.25, Other 2.21±0.70. **(C)** Merged confocal micrograph of SSp-L6b^Ctgf^ (tdT, red) neurobiotin filled (green) recorded neurons in a brain coronal slice. Lower panel depicts example reconstructions of three neurobiotin filled L6b^Ctgf^ neurons. **(D) Spontaneous activity of SSp-L6b^Ctgf^ neurons in whole-cell patch-clamp mode.** Representative current-clamp whole-cell recordings of the membrane potential dynamics of silent (top) and spontaneously active (bottom) L6b^Ctgf^ neurons. Boxed region shown on an expanded scale below, highlighting heterogeneous firing patterns, including bursting activity. **(E) Distribution of response types.** Proportion of spontaneously active (14/30) and silent (16/30) L6b^Ctgf^ neurons under baseline conditions. Lower scatter-plot shows that active L6b^Ctgf^ neurons fire spontaneous APs at a mean frequency of 4.4±1.6Hz (14 cells). **(F) Passive membrane properties of SSp-L6b^Ctgf^ neurons in whole-cell patch-clamp mode.** Resting membrane potential was significantly more depolarized in active L6b^Ctgf^ (14 cells) neurons (*vs.* silent 16 cells). Note, membrane potential was corrected for the liquid junction potential +16mV. Cell capacitance and membrane resistance did not differ between groups (15 active and 14 silent). Passive properties were measured after membrane rupture and stabilization of the whole-cell configuration. Stats for resting membrane potential: unpaired t test active *vs.* silent L6b^Ctgf^ t(28)=2.415, p=0.023. Mean±SEM (mV): active-85.0±1.5, silent-88.8±0.7. Stats for cell capacitance: unpaired t-test active *vs.* silent L6b^Ctgf^ t(27)=0.3986, p=0.69. Mean±SEM (pF): active 68±5, silent 72±6. Stats for membrane resistance: Mann-Whitney active *vs.* silent L6b^Ctgf^ U=85, p=0.39. Mean±SEM (MΩ): active 270±55, silent 206±26. **(G)** Input resistance plotted as a function of resting membrane potential. Input resistance was calculated from the steady-state membrane voltage response to a-100pA current injection and did not correlate with membrane potential. Data are shown as individual mean values. (Pearsońs r=0.28, p=0.14, 29 cells). **(H)** Representative trace of a L6b^Ctgf^ neuron’s automated detection of action potentials (blue asterisks). Right, probability distribution of the inter spike interval (ISI) for all detected events. A threshold for multispike detection was set at the interstimulus interval between the two distribution peaks (43.5ms, dashed line). **(I)** Examples of diverse firing patterns observed in L6b^Ctgf^ active neurons, including doublets interspersed with single spikes (left) and high-frequency bursts (right). **(J)** Quantification of the average number of spikes per multispike event (left) and their average ISI (right). No differences were found between L6b^Ctgf^ and L6b^Other^ neurons. Stats for spike sequence: Mann-Whitney U=9, p=0.076. Mean±SEM (n): L6b^Ctgf^ 3.86±1.71, L6b^Others^ 2.72±0.26. Median: L6b^Ctgf^ 2.0, L6b^Others^ 2.83. Hodges-Lehmann median difference 0.33. Stats for doublets inter-spike interval: Mann-Whitney U=5, p=0.17. Mean±SEM (ms): L6b^Ctgf^ 27.4±3.15, L6b^Others^ 19.2±5.48. Median: L6b^Ctgf^ 29.1, L6b^Others^ 16.5. Hodges-Lehmann median difference-7.94. SSp, primary somatosensory cortex; MOp, primary motor cortex. Panels A-B related to **Figure 4**. Panels C-J related to **Figure 5**. For statistical details see **Table S3**.

**Supplementary Figure 3.**
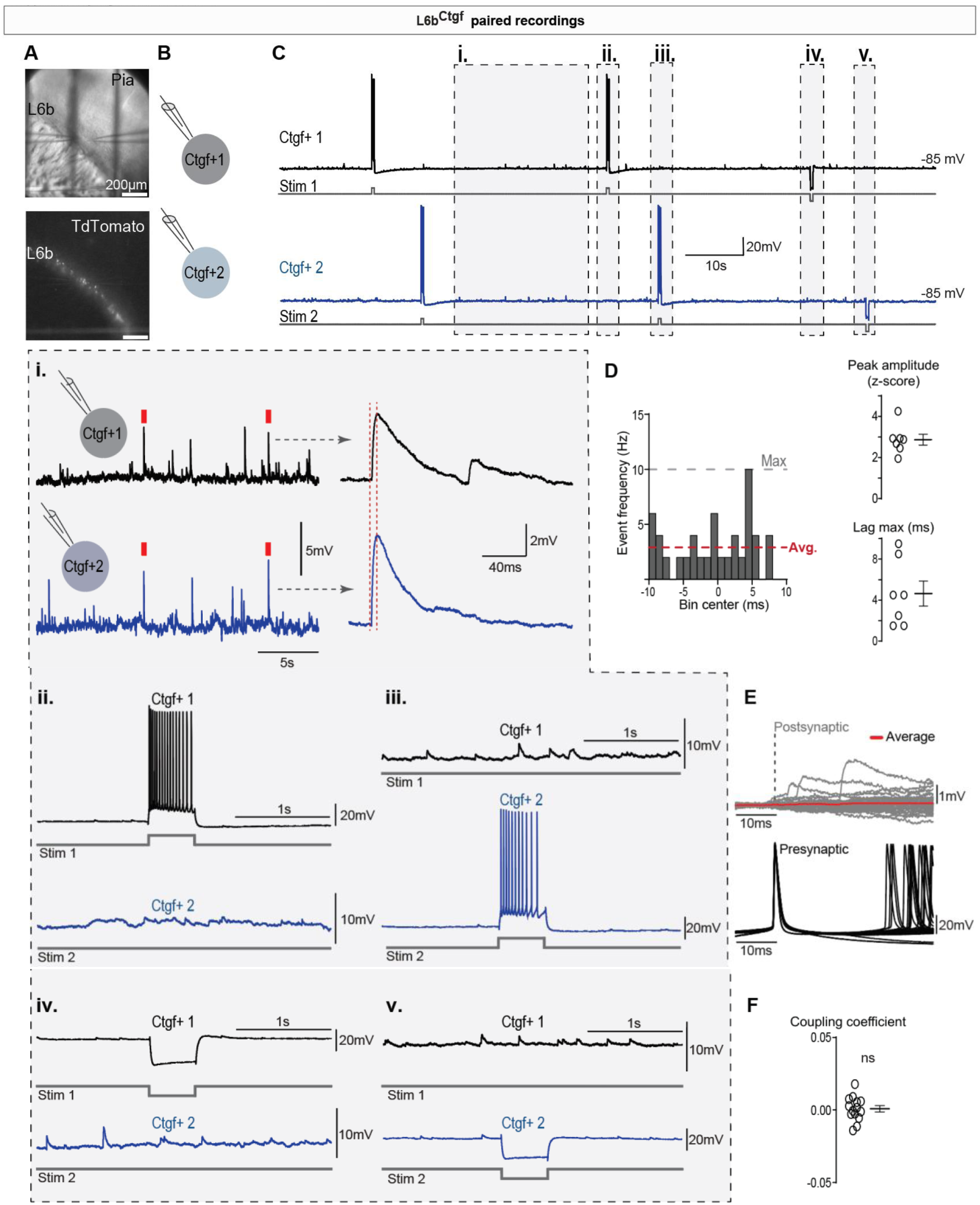
Patched pairs of SSp-L6b^Ctgf^ neurons are neither synaptically nor electrically coupled. **(A)** Microphotograph of a cortical slice in a coronal plane with patch electrodes (top, differential interference contrast image) and corresponding fluorescent image of L6b^Ctgf^ neurons (bottom, tdT). **(B)** Schematic illustration of a paired patch-clamp of L6b^Ctgf^ neurons. **(C)** Representative simultaneous patch-clamp recordings in current-clamp mode from a pair of L6b^Ctgf^ neurons. The black trace represents the membrane voltage recorded from L6b^Ctgf^ neuron 1 (Ctgf^+^1), and the blue trace from L6b^Ctgf^ neuron 2 (Ctgf^+^2). The stimulus protocol for each neuron is illustrated below the trace (Stim 1 and Stim 2). Segments highlighted with boxes are depicted on an expanded scale bellow. **(i)** Spontaneous post-synaptic potentials (PSPs) recorded under basal conditions. Examples of simultaneously occurring events are marked (red box). Right, events depicted on the expanded scale with red dashed lines indicating the onset and peak of PSPs in Ctgf^+^1 and Ctgf^+^2. **(ii-iii)** Evoked trains of APs in one neuron (current step +100pA, 500ms) did not elicit responses in the paired neuron (14 directions, seven pairs, five mice). **(iv-v)** Membrane potential hyperpolarization in one neuron (current step - 100pA, 500ms) did not elicit responses in the paired neuron. **(D)** Spontaneous PSP-time cross-correlogram from an example L6b^Ctgf^ neuron pair. Gray dashed line indicates a cross-correlation maximum. Red dashed line indicates the average cross-correlation value. Summary plot for spontaneous PSPs peak amplitude (z-score) and latency of the cross-correlation maximum (Lag max) for all pairs. Stats for synchrony: Mean±SEM: 2.9±0.3 z-score and Lag max 4.6±1.2ms. Seven pairs, five mice. **(E)** Evoked APs in Ctgf^+^1 did not induce time-locked PSPs in Ctgf^+^2. Example APs (3 trains, 500ms positive-current steps) in Ctgf^+^1 (black, below) and corresponding membrane potential in Ctgf^+^2 (gray, above). Red trace indicates the average membrane potential of Ctgf^+^2. **(F)** Coupling coefficient (CC) of L6b^Ctgf^ neuron pairs. CC was determined as the ratio between Ctgf^+^2 and Ctgf^+^1 junctional steady state voltage deflection; measured during injection of a hyperpolarizing current step into Ctgf^+^1, as shown in **C.iv-v**. Stats for CC: one-sample t-test *vs.* zero t(13)=0.3859, p=0.71. Mean±SEM 0.00086±0.00222. 14 directions, seven pairs, five mice.

**Supplementary Figure 4.**
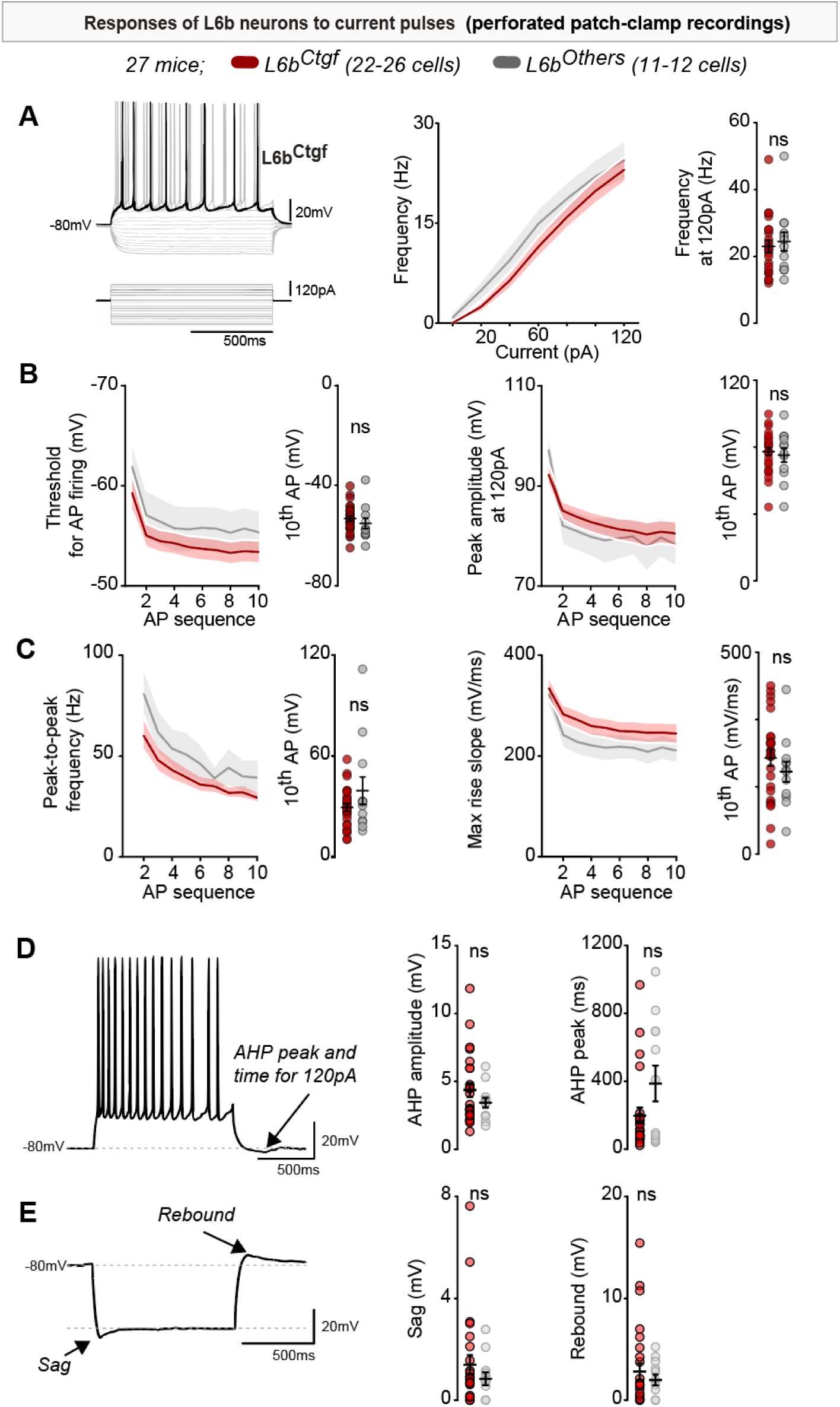
L6b^Ctgf^ and L6b^Others^ neurons show similar active properties. **(A)** Left, representative trace from a L6b^Ctgf^ neuron responding to current pulses. Middle, input-output curves of L6b^Ctgf^ neurons (red) and L6b^Other^ neurons (black). Right, firing frequency in response to depolarizing current steps. Stats for firing frequency response to 120pA: Mann-Whitney U=120.5, p=0.69. Mean±SEM (Hz): L6b^Ctgf^ 23.0±1.86, L6b^Others^ 24.4±2.85. Median: L6b^Ctgf^ 22.0, L6b^Others^ 24.0. Hodges-Lehmann median difference 1.0. **(B)** No differences were found between L6b^Ctgf^ and L6b^Others^ neurons in AP threshold (left) or peak amplitude (right). Stats for AP firing threshold: Mann-Whitney U=102, p=0.18. Mean±SEM (mV): L6b^Ctgf^-53.4±1.09, L6b^Others^-55.4±2.16. Median: L6b^Ctgf^-53.9, L6b^Others^-57.5. Hodges-Lehmann median difference-2.81. Stats for Peak AP amplitude: Unpaired t-test t(36)=0.47, p=0.64. Mean±SEM (mV): L6b^Ctgf^ 80.6±2.4, L6b^Others^ 78.4±4.3. **(C)** No differences in peak-to-peak frequency (left) or max rise slope (right). Stats for Peak-to-peak frequency: Mann-Whitney U=139.0, p=0.60. Mean±SEM (Hz): L6b^Ctgf^ 29.4±2.34, L6b^Others^ 39.4±8.15. Median: L6b^Ctgf^ 29.1, L6b^Others^ 31.5. Hodges-Lehmann median difference 2.33. Stats for Max rise slope: Unpaired t-test t(36)=0.99, p=0.33. Mean±SEM (mV/ms): L6b^Ctgf^ 244.9±20.4, L6b^Others^ 210.6±25.1. **(D)** Left, representative trace of a L6b^Ctgf^ neuron, after-hyperpolarization (AHP) is measured as the membrane potential lowest point at the end of the current pulse (arrow). Stats for AHP amplitude: Mann-Whitney U=130.0, p=0.53. Mean±SEM (mV): L6b^Ctgf^ 4.37±0.51, L6b^Others^ 3.44±0.36. Median: L6b^Ctgf^ 3.30, L6b^Others^ 3.39. Range: L6b^Ctgf^ [1.31 to 11.84], L6b^Others^ [1.74 to 6.10]. Hodges-Lehmann median difference-0.48. Stats for Time to reach trough of after-hyperpolarization: Mann-Whitney U=128.0, p=0.49. Mean±SEM (ms): L6b^Ctgf^ 198.4±46.2, L6b^Others^ 387.1±105.6. Median: L6b^Ctgf^ 112.0, L6b^Others^ 253. Range: L6b^Ctgf^ [23.0 to 969.0], L6b^Others^ [41.0 to 1046]. Hodges-Lehmann median difference 27.0. **(E)** Left, representative trace of a L6b^Ctgf^ neuron in response to a-120pA current, arrow 1 (Sag), arrow 2 (rebound potential). Note, the sag was calculated in response to a-120pA hyperpolarizing step and rebound was calculated from-180pA hyperpolarizing step. Right, no differences were found for Sag or rebound amplitude. Stats for Sag: Mann-Whitney U=125.5, p=0.54. Mean±SEM (mV): L6b^Ctgf^ 1.39±0.38, L6b^Others^ 0.84±0.25. Median: L6b^Ctgf^ 0.81, L6b^Others^ 0.75. Hodges-Lehmann median difference-0.12. Stats for Rebound amplitude: Mann-Whitney U=153.0, p=0.94. Mean±SEM (mV): L6b^Ctgf^ 2.80±0.82, L6b^Others^ 1.98±0.54. Median: L6b^Ctgf^ 0.85, L6b^Others^ 1.7. Hodges-Lehmann median difference 0. (A-E) L6b^Ctgf^ 22-26 cells. L6b^Others^ 11-12 cells. 27 mice. Related to figure **Figure 5** and **Figure S2H-J**. For statistical details see **Table S3**.

**Supplementary Figure 5.**
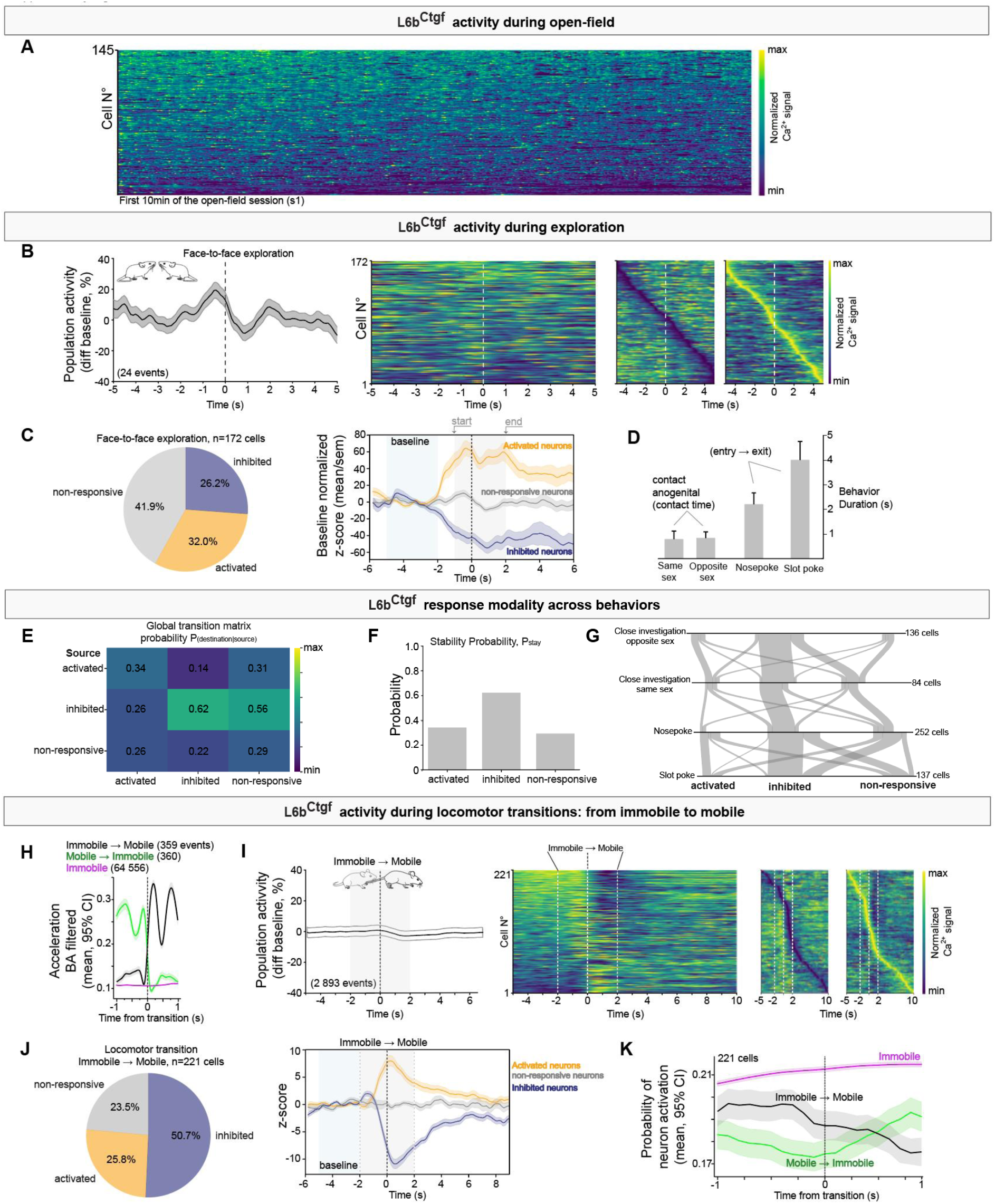
In vivo calcium imaging of L6b^Ctgf^ neurons in freely moving mice. **(A)** Calcium dynamics of 145 L6b^Ctgf^ neurons during the first 10min of open-field exploration during session 1 (empty chamber, four mice). Sorted according to min-max activity. Time is binned in 1s intervals. **(B)** Face-to-face social exploration (four mice). Left, grand average fluorescence signal, across all recorded neurons, around the annotated behavioral timepoint (vertical dashed line: contact onset). Heatmaps showing normalized cell activity averaged across behavioral events, organized by activity level; most active on top (left), time of peak (middle) or time of trough (right). **(C)** Left, the proportion of cells that are classified as inhibited (blue), activated (orange) or non-responsive (gray) during the behavioral window. Right, behavior aligned average peri-stimulus time histograms (PSTHs) for each cell class. Period used for baseline z-score correction (light blue shade) and window of time used for response class identification (light gray shade). Lines represent the baseline normalized class response, z-score mean±SEM. **(D)** Bar graph showing the mean bout duration for exploratory behaviors in which the population-averaged signal exhibited a decreased response. **(E)** Global transition probability matrix showing the likelihood that a cell classified as activated, inhibited, or non-responder during one behavior retains or changes its classification during another behavior. Probabilities are row-normalized and pooled across all pairwise behavior comparisons. **(F)** Stability probability per class (Pstay), defined as the diagonal of the global transition matrix (E), indicating the likelihood that cells preserve their response classification across behaviors. **(G)** Alluvial plot of response class stability and switching. For each behavior, the proportion of cells that maintain or switch response class is shown. Line thickness is proportional to the fraction of cells in each transition. Vertical lines indicate class stability, whereas crossing lines indicate class switching. **(H)** Head-mounted accelerometer signals during locomotor state transitions. Acceleration magnitude derived from a head-mounted accelerometer using the body acceleration (BA) filter. Traces show two locomotor transition windows (1s before and 1s after transition detection) and a reference period of sustained immobility (2s). **(i)** Locomotor initiation. Transition windows were defined as 2s of detected immobility followed by 2s of mobility (gray shading). The dashed line marks the moment of locomotor initiation. Population grand-average fluorescence traces and corresponding heat maps are shown as in (B). **(j)** Locomotor initiation. By class proportions and normalized z-score as in (C). **(K)** Peri-transition L6b activation probability around locomotor state changes. Population averaged probability that L6b^Ctgf^ neurons are active, aligned to the behavioral transition (t=0s; dashed line). Traces show transitions from immobile-to-mobile (black) and mobile-to-immobile (green), with an immobility reference condition (immobile, remains immobile; pink). Shaded regions indicate 95% confidence intervals across events. Calculated for all events shown in (H). Related to figure **Figure 6** and **Figure 7**.

## SUPPLEMENTARY TABLES

- **Table S1.** Number of animals, and samples analyzed, per experiment. Related to Figures 1-7.
- **Table S2.** Inclusion and exclusion criteria per experiment. Related to Figures 1-7.
- **Table S3.** Statistical analysis. Related to Figures 4 and Figure 5, and Supplementary Figures 2 and 4. Statistical tests used for all experimental comparisons. Numbers in parentheses indicate sample size (n). Data are reported as mean±SEM.
- **Table S4.** Acronyms for anatomical areas. Related to Figure 3. Acronyms are listed by panel in the order shown. Corresponding macro-regions are provided where applicable.

***------- File END ------***

## Notes

https://github.com/DM-DBB/L6b_CTGF_project.git

https://doi.org/10.5281/zenodo.19002258

### Competing Interest Statement

The authors have declared no competing interest.

