## Supplementary material for "Cortical Layer 6b Persistent Subplate Neurons Reciprocally Connect Sensorimotor Areas and Inversely Reflect Somatosensory Engagement": Table S1

### Table S1: Number of animals and samples analyzed per experiment. Related to Figures 1-7.

| Experiment | Animals (N) | Samples / Cells (n) |
| --- | --- | --- |
| RNAscope and anatomy |  |  |
| RNAscope | 6 | 2 slides |
| Coronal batch tracing | SSp (4), MOp (4) | 91 samples |
| Cell counts from sagittal sections | SSp (2), MOp (2) | L6b^Ctgf^ total n = 132,891  L6b^Ctgf^ cortical n = 128,291  L6b^Ctgf^ non-cortical n = 4,589 |
| Optogenetics |  |  |
| Spike fidelity in Ctgf+ neurons | 14 | 5 cells |
| SSp responders |  | Pyr (19), FSI (11), others (7) |
| SSp nonresponders |  | Pyr (4), FSI (5), others (3) |
| MOp responders |  | Pyr (10), FSI (5), others (7) |
| MOp nonresponders |  | Pyr (2), FSI (0), others (2) |
| Paired pulse ratio (SSp) |  | Pyr (19), FSI (11), others (7) |
| Paired pulse ratio (MOp) |  | Pyr (10), FSI (5), others (7) |
| Perforated patch characterization | | |
| Baseline intrinsic properties | 29 | L6b^Ctgf^ (33), L6b^Other^ (18) |
| Cells receiving current steps |  | L6b^Ctgf^ (22–26), L6b^Other^ (11-12) |
| MATLAB multispiking analysis |  | L6b^Ctgf^ (17), L6b^Other^ (8) |
| Multispiking vs tonic firing |  | L6b^Ctgf^ multispiking (9), tonic (8); L6b^Other^ multispiking (5), tonic (3) |
| Population activity |  |  |
| In vivo Ca2+ imaging | 4 | 255 neurons |

N indicates number of animals; n indicates number of cells or samples analyzed.
