## Supplementary material for "Cortical Layer 6b Persistent Subplate Neurons Reciprocally Connect Sensorimotor Areas and Inversely Reflect Somatosensory Engagement": Table S2

### Table S2. Inclusion and exclusion criteria per experiment. Related to Figures 1-7.

| Experiment / Step | **Inclusion criteria** | **Exclusion criteria** |
| --- | --- | --- |
| RNAScope |  |  |
| Genotype | *Ctgf-T2A-dgCre* | N/A |
| Age / sex | P59+, mixed sex | N/A |
| Tissue collection | Fresh frozen brain collection | N/A |
| Sectioning | Cryostat, every 5th section collected | Damage to section during cutting |
| Sampling | 2 slides at same somatosensory cortex level | N/A |
| Signal quality | Distinct fluorescent puncta | Overlapping puncta across channels |
| Nuclear integrity | DAPI signal with intact nuclear morphology | Smeared DAPI signal |
| Imaging | 10X overview; 40X images analyzed | N/A |
| Quantification (manual) | Distinct puncta clusters around DAPI nuclei counted as cells | N/A |
| Quantification (scripted) | Centroid-based segmentation; allowance for size=80-350 circularity=0.30-1.00 | Cells outside threshold range |
| Anatomy |  |  |
| Genotype | *Ctgf-T2A-dgCre* (coronal); *Ctgf-T2A-dgCre; tdTomato* (sagittal) | N/A |
| Age / sex | P60+, mixed sex | N/A |
| Surgery | Unilateral stereotaxic virus injection (GFP + mRuby) | N/A |
| Incubation | ≥4 weeks post-injection | N/A |
| Fixation | Perfusion; overnight PFA fixation; PBS rinse | N/A |
| Sample embedding | 4% agarose | N/A |
| Sectioning | Vibratome; all slices collected consecutively | Slice damaged during cutting |
| Staining | GFP & DsRed & Neurotrace (coronal); GFP (sagittal) | N/A |
| Mounting | ProLong antifade, coverslip #1.5, no bends | N/A |
| Imaging signal quality | Clear GFP+ fibers prior to counterstaining | No GFP+ fibers could be seen prior to counterstaining |
| Mapping / analysis | Injection site shows visible GFP+ cell bodies | Visible GFP+ cell bodies in other brain areas but the one injected |
| Sampling consistency | At least four slices representing each distinct area/per mouse | N/A |
| Optogenetics –Electrophysiology | | |
| Animals | *Ctgf-T2A-dgCre* | N/A |
| Age / sex | P60+, mixed sex | N/A |
| Surgery | Unilateral stereotaxic virus injection (GFP + mRuby) | N/A |
| Incubation | ≥4 weeks post-injection | N/A |
| Fixation | Perfusion; overnight PFA fixation; PBS rinse | N/A |
| Viral expression | Positive fluorescence | N/A |
| Current clamp | Action potentials evoked by current steps | No action potentials |
| Voltage clamp | Detectable synaptic inputs | No inputs in trace |
| Responder definition | Time-locked responses to light | N/A |
| Non-responder definition |  | No time-locked responses |
| Statistical analysis | All responders and nonresponders counted | N/A |
| Pharmacology / PPR | Responders completing full protocol | Nonresponders excluded |
| PPR analysis | All responder cells under gabazine | N/A |
| Perforated patch |  |  |
| Genotype | *Ctgf-T2A-dgCre; tdTomato* | N/A |
| Age / sex | P28-P35, mixed sex | N/A |
| Treatment | Trimethoprim (TMP) injection | N/A |
| Electrophysiology coronal slice thickness | 250µm | N/A |
| Region | SSp (bregma 0.50–0.02 mm) | N/A |
| Target confirmation | L6b and white matter fibers under DIC/tdTomato | N/A |
| Cell shape | Elongated or pyramidal soma; visible dendrites | N/A |
| Nucleus | Visible nucleus; sharp contours | N/A |
| Electrical seal | GΩ seal held ≥10s | N/A |
| Series resistance | Ra <100 MΩ after 15-20min | Ra change >20% |
| Baseline stability | <1mV drift over 1min | N/A |
| AP amplitude | >70mV | Failure to fire APs |
| Statistical analysis | All cells meeting criteria counted | Unstable baselines excluded |
| Passive properties | RMP and capacitance reported for all included cells | N/A |
| MATLAB analysis | Active cells with ≥5 min baseline | Baselines <5 min excluded |
| Evoked activity | Cells receiving -120, -180, and +120pA steps | N/A |
| In vivo Ca2+ imaging |  |  |
| Genotype | *Ctgf-T2A-dgCre* | N/A |
| Age | 2-5 months, mixed sex | N/A |
| Trace selection for analysis | PCA-based selection | N/A |
| Quality control |  | Photobleaching after 40 min |
| Normalization | Longitudinal projection processing | N/A |

Criteria were applied consistently across experiments.
